# Level of host concealment shape parasitoid community of microlepidopteran species living on hops

**DOI:** 10.1101/2025.04.06.647420

**Authors:** Tomáš Hovorka, Kamil Holý, Cristina Vasilita, Lars Krogmann, Petr Janšta

**Author notes:** **Corresponding author:** Tomáš Hovorka. **Data availability statement:** Data used in this manuscript are available from the Zenodo open repository https://doi.org/10.5281/zenodo.15119124.

## Abstract

**Background:** Parasitoid-host interactions are key drivers of insect community structure, with host concealment influencing parasitoid diversity and parasitism rates. However, the effectiveness of different host defense strategies against parasitoids remains insufficiently understood.

**Objective:** This study examines how host concealment level affects parasitoid communities and parasitism rates in two microlepidopteran species developing on hops (*Humulus lupulus* L.), *Caloptilia fidella* and *Cosmopterix zieglerella*, which employ leaf-rolling and leaf-mining strategies, respectively.

**Methods:** We combined morphological identification with molecular species delimitation using ITS2 and CO1 markers and applied ASAP and bPTP methods to refine parasitoid taxonomy and detect cryptic species.

**Results:** Semi-concealed *C. fidella* larvae in leaf rolls experienced significantly higher parasitism than their mining stages, whereas fully concealed *C. zieglerella* had lower parasitism rates. Molecular analyses confirmed the idiobiont strategy in *Sympiesis acalle*, *S. sericeicornis*, and *Elachertus fenestratus*, and bPTP proved more sensitive in detecting cryptic species than ASAP.

**Significance:** These findings demonstrate that semi-concealed hosts face a higher risk of parasitism than fully concealed hosts, suggesting that leaf-mining provides better protection than leaf-rolling in studied hosts. The study also highlights the power of molecular tools in species delimitation, emphasizing their importance for refining parasitoid taxonomy and advancing our understanding of host-parasitoid interactions.

## INTRODUCTION

Insect herbivores and their natural enemies, including many parasitoids, constitute a substantial part of global insect diversity and suggests plant-herbivore-parasitoid relationships as one of the major components of terrestrial ecosystems (Hawkins, 1994; Quicke, 1997; Price, 2002; Sugiura, 2007; Smith et al., 2008; Hamilton et al., 2010). Currently, more than half of all known terrestrial species are part of plant-herbivore-parasitoid food webs (Mey, 1996; Smith et al., 2008; Novotný et al., 2010, Leppänen et al., 2013). These food webs include both organisms providing ecosystem services such as pollinators and natural enemies of various organisms, as well as economically important pests in agriculture and forestry (Kremen & Chaplin-Kramer, 2007; Letourneau et al., 2009; Martin et al., 2013; Gonzales et al., 2022). The interactions of these organisms within food webs have led herbivore hosts to evolve a plethora of various defence strategies (Brodie & Smatresk, 1990; Stankowich & Campbell, 2016; Aoyama & Ohshima, 2019).

A common primary defence strategy of herbivores against enemies such as parasitoids is the creation of physical barriers. Such a strategy is known from a large number of small arthropods, living inside plant material, which, apart from food, serves as protective physical barrier against predators and parasitoids (Aoyama & Ohshima, 2019). The ability of insects to use plants as their shelter by creating various structures (leaf rolls, galls, mines) is a key factor influencing the parasitoid community (Hering, 1951; Fernandes & Price, 1988; Sinclair & Hughes, 2010, Hrček et al., 2013). A partially hidden lifestyle (e.g. leaf rolls) as a defence strategy against parasitoids appears to be the least advantageous strategy compared to living on the surface of plants (Hrček et al., 2013), where individuals are more frequently attacked by predators (Libra et al., 2019). Hawkins (1994) and Hrček et al. (2013) stated that parasitism rate and parasitoid species richness increase with the degree of concealment from exposed feeders through leaf rollers, leaf tiers and case-bearers, reach maximum values in leaf miners and gallers, and then decrease again in even more concealed borers and root-feeders. In contrast, other studies have shown that exposed hosts can exhibit high parasitism rates (e.g. Sarfraz et al., 2005; Konvičková et al., 2024), and that other factors, such as seasonality, may be more important than host concealment (Le Corff et al., 2000). To address this issue, it is necessary to obtain and analyse additional data from various locations around the world.

Despite that, life inside the plant tissue (e.g. leaves) and creating mines is one of the most common life strategies of phytophagous insects (Tooker & Giron, 2020). Even if mining is not a perfect strategy against parasitoids, mines and specifically their shapes have been suggested as a defence mechanism to avoid or confuse parasitoids (Needham et al., 1928; Connor & Taverner, 1997; Aoyama & Ohshima, 2019). During larval development, some species of mining butterflies (e.g. *Acrocercops transecta* Meyrick, 1931) create variously complex and shaped corridors inside plant leaves and increase the area over which the parasitoid must search for its host or provide a space to escape from the parasitoid in case the host is found (Ayabe et al., 2008; Aoyama & Ohshima, 2019). As the complexity of mines increases, their level of parasitism decreases. Thus, the complexity of mines is partly a result of the prey-predator relationship between mining species and their parasitoids. For example, the mining moth *Phyllonorycter malella* (Gerasimov, 1931) (Lepidoptera: Gracillaridae) creates a complex network of mines with unconsumed tissue, which serves as protection (Djemai et al., 2000). The more complex the structure of the mines in the leaf of this species is, the less the caterpillar was parasitized (Aoyama & Ohshima, 2019). However, the exact role of mining as a defence mechanism against parasitoids is still disputable. Several studies (i.e. Askew, 1980; Hawkins & Lawton, 1987; Hawkins, 1990, 1994) have shown mining insect species to have more parasitoids than hosts from any other feeding guild (Connor & Taverner, 1997). However, mining still may be one of the initial strategies to avoid parasitoids. The taxonomic composition of the parasitoid community attacking leaf-mining sawflies (Hymenoptera: Tenthredinidae) has been shown as radically different from that attacking external-feeding sawflies having less species of braconid and ichneumonid (both Hymenoptera: Ichneumonoidea) parasitoids (Pschorn-Walcher & Altenhofer, 1989). With the decrease in the number of braconid and ichneumonid parasitoids, there was an increase in the number of species from the family Eulophidae (Hymenoptera: Chalcidoidea), which generally parasitize concealed-feeding Lepidoptera, such as leaf miners and leaf rollers (Stireman & Shaw, 2022). This aligns well with observations by Pschorn-Walcher and Altenhofer (1989), who suggested that the initial escape from parasitism associated with adopting a concealed-feeding habit, such as leaf mining or rolling, may have provided sufficient impetus to selectively reinforce these feeding strategies. These adaptations likely led to interactions with a distinct and potentially more specialized group of parasitoids, as evidenced by studies like Hrček et al. (2013).

Comparing the parasitism rates and the success of different feeding strategies, such as leaf-mining and leaf-rolling, provides a deeper understanding of how these strategies function as defences against parasitoids. Some microlepidopteran species rely exclusively on the mining strategy, while others adopt a combination of mining and rolling during their larval stages (Tooker & Giron, 2020). This raises the question of how these two strategies are comparable in terms of effectiveness against parasitoid attacks. Therefore, it is essential to evaluate the benefits and limitations of each strategy to better understand their evolutionary significance and ecological implications.

In this study we analysed the parasitoid community of two microlepidopteran species on hops (*Humulus lupulus* L.), *Caloptilia fidella* (Reutti, 1853) (Lepidoptera: Gracillaridae) and *Cosmopterix zieglerella* (Hübner, 1810) (Lepidoptera: Cosmopterigidae), with similar mine complexity but different life strategies through the larval life cycle.

With a wingspan of 9–12 mm, *Caloptilia fidella* is one of the smallest West-Palaearctic moths in the genus *Caloptilia* Hübner, 1825. This species is bivoltine, with larvae appearing mainly in July and September, while adults overwinter. The larvae primarily feed on hop but occasionally also on *Celtis australis*. During development, the larvae first create a triangular leaf mine (1st-3rd instar) before moving to feed externally within a rolled leaf tip or lobe (4th-5th instar). Pupation occurs in silken cocoons on the underside of the host plant’s leaves (Figure 3A-D; Baugnée & De Prins, 2010; Laštůvka et al., 2018; Watson et al., 2021).

The moth *Cosmopterix zieglerella* (a wingspan measuring 8–11 mm) is distributed throughout the Palaearctis. This species is monovoltine, with larval activity occurring from July to September. The larvae hibernate within a silken cocoon located in ground detritus. They exhibit monophagy, feeding exclusively on hop. During the initial stages, the larvae construct a narrow, irregular linear mine along the leaf vein, lined with silk and serving as a protective shelter. The larvae of *C. zieglerella* are green during the early instars of their development. Prior to pupation, the larva acquires a distinctive coloration characterized by red stripes. As development progresses, the mine expands into a diffuse, irregular yellowish-white blotch containing scattered frass (Figure 3E-H; Koster & Sinev, 2003; Laštůvka et al., 2018).

This study investigates the interactions between two host lepidopteran species *C. fidella*, *C. zieglerella*, and their associated parasitoids, focusing on parasitoid species composition, bionomy, and parasitism rates. The aims are to (i) reveal how the contrasting defensive strategies of the two hosts affect their susceptibility to parasitism, (ii) identify key parasitoid species and their life histories, and (iii) assess ecological factors shaping host-parasitoid dynamics.

## MATERIAL AND METHODS

### Sampling and rearing

Sampling was conducted between 2020 and 2022 across 18 sites in the Czech Republic, Slovakia, Hungary, Romania, and Croatia (Table 1). Whole hop leaves containing 1^st^ or 2^nd^ generation larvae of *C. fidella* (1st-5th instar) were collected, with a total of 50 leaves per site. For *C. zieglerella*, leaves with visible mines were collected in as many numbers as available per site (always < 50 leaves per location) where the species was present. Collected leaves containing mines or rolls with host species were transported to the laboratory in separate zip-lock bags for each host and site. Upon arrival, each leaf was carefully inspected and cleaned of any other insect hosts, most commonly (aphids, spider mites, planthopper nymphs and whiteflies). A sufficiently large section of the leaf containing the mine (*C. zieglerella*) or the corresponding host stage (*C. fidella*) was then clipped to allow the host species or its parasitoid to complete its life cycle and placed in sterile plastic petri dish (100 mm diameter, 15 mm height). The petri dishes were stored under controlled conditions (20°C, 75% humidity, and a 16:8 h light:dark cycle) in Trigon Plus ST 2 B SMART climate chambers (TRIGON PLUS Ltd., Czech Republic) for 40 days. Every two days until adult insects emerged, the petri dishes were inspected, and the leaves were moistened with a water solution of the fungicide LUNA® (fluopyram 200 g/L and tebuconazole 200 g/L; Bayer AG, Germany) using a laboratory sprayer to prevent mould during rearing. This fungicide has no adverse effects on arthropods (tested by Central Institute for Supervising and Testing in Agriculture, Czech Republic). The specific developmental biology of the host and its parasitoids was recorded. The biology of the parasitoids was determined during their development on the hosts using forceps and dissecting needles to carefully open the leaf shelters, ensuring minimal disruption to their development. Additionally, parasitoids were monitored to determine whether they were koinobionts or idiobionts and classified as either ectoparasitoids or endoparasitoids. Whenever feasible, individual parasitoids were photographed. Direct observation of parasitoid life strategies was supplemented by post-emergence dissection of leaf mines and shelters to analyze remnants of host larvae, pupae, parasitoid cocoons, and the positions of emergence or exit holes. Additionally, the pupae or cocoons of parasitoids were monitored for the potential occurrence of hyperparasitoids. After emergence, the adults of hosts or parasitoids were stored in 96% ethanol at −20°C until DNA isolation.

**Table 1.**
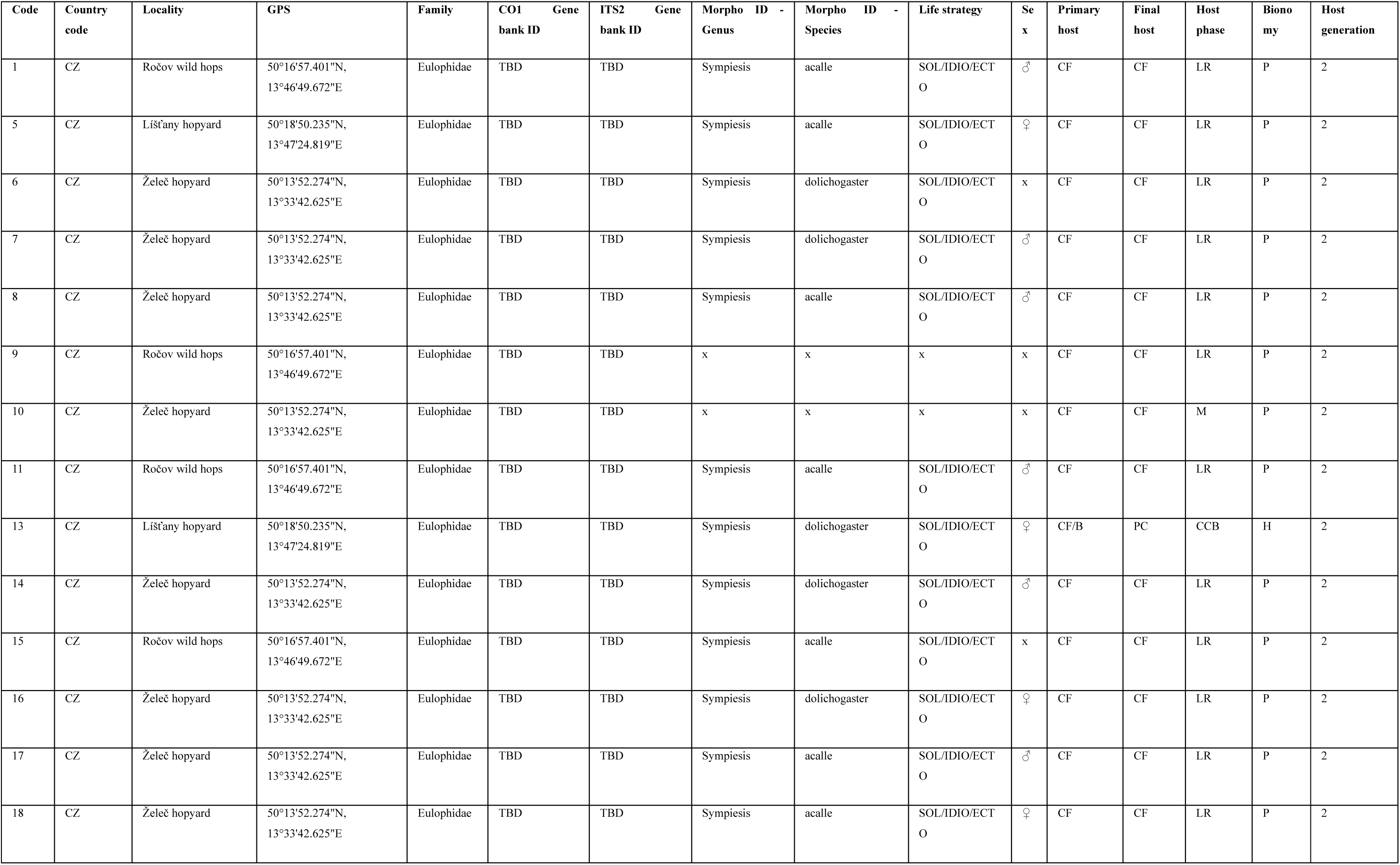
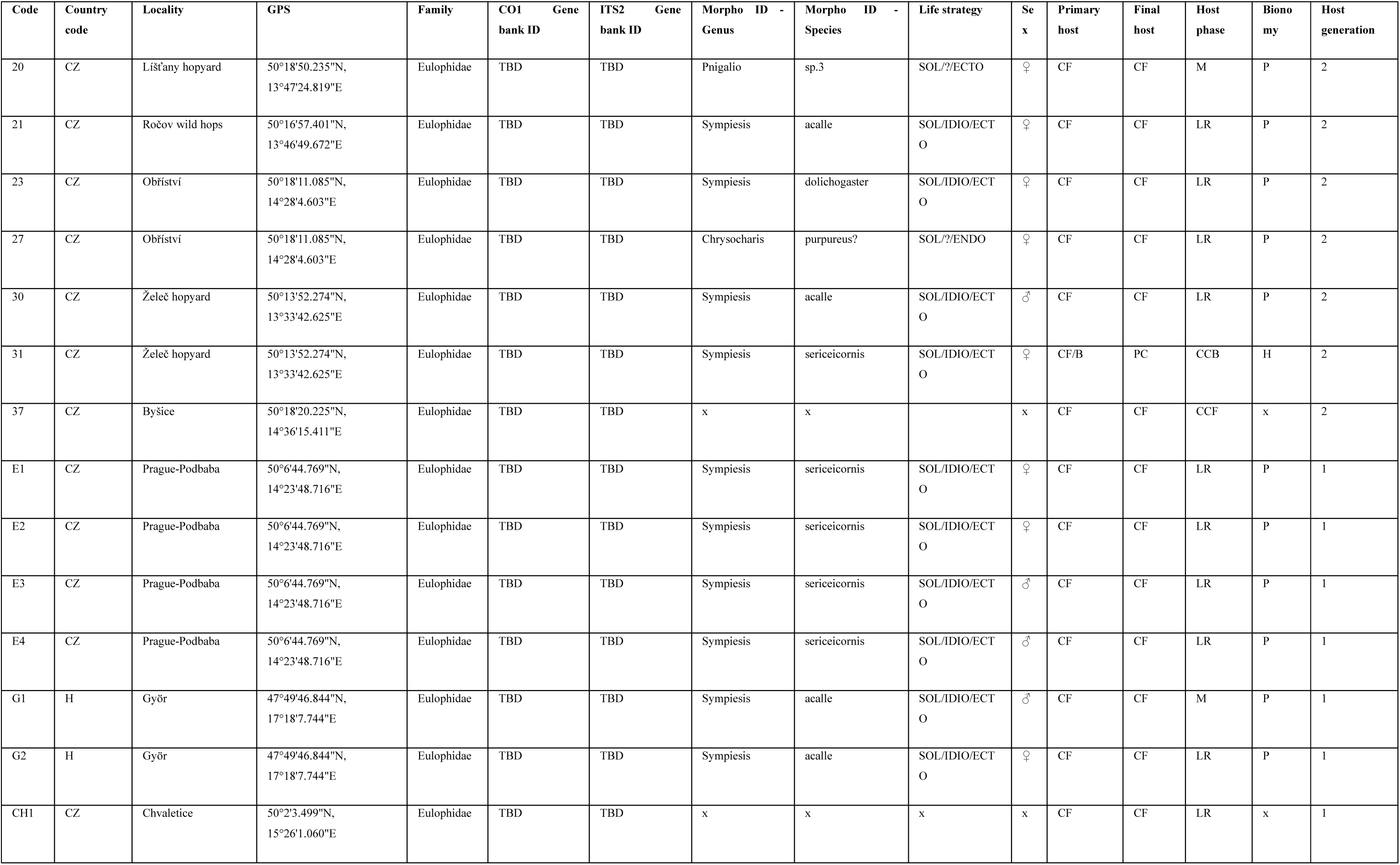
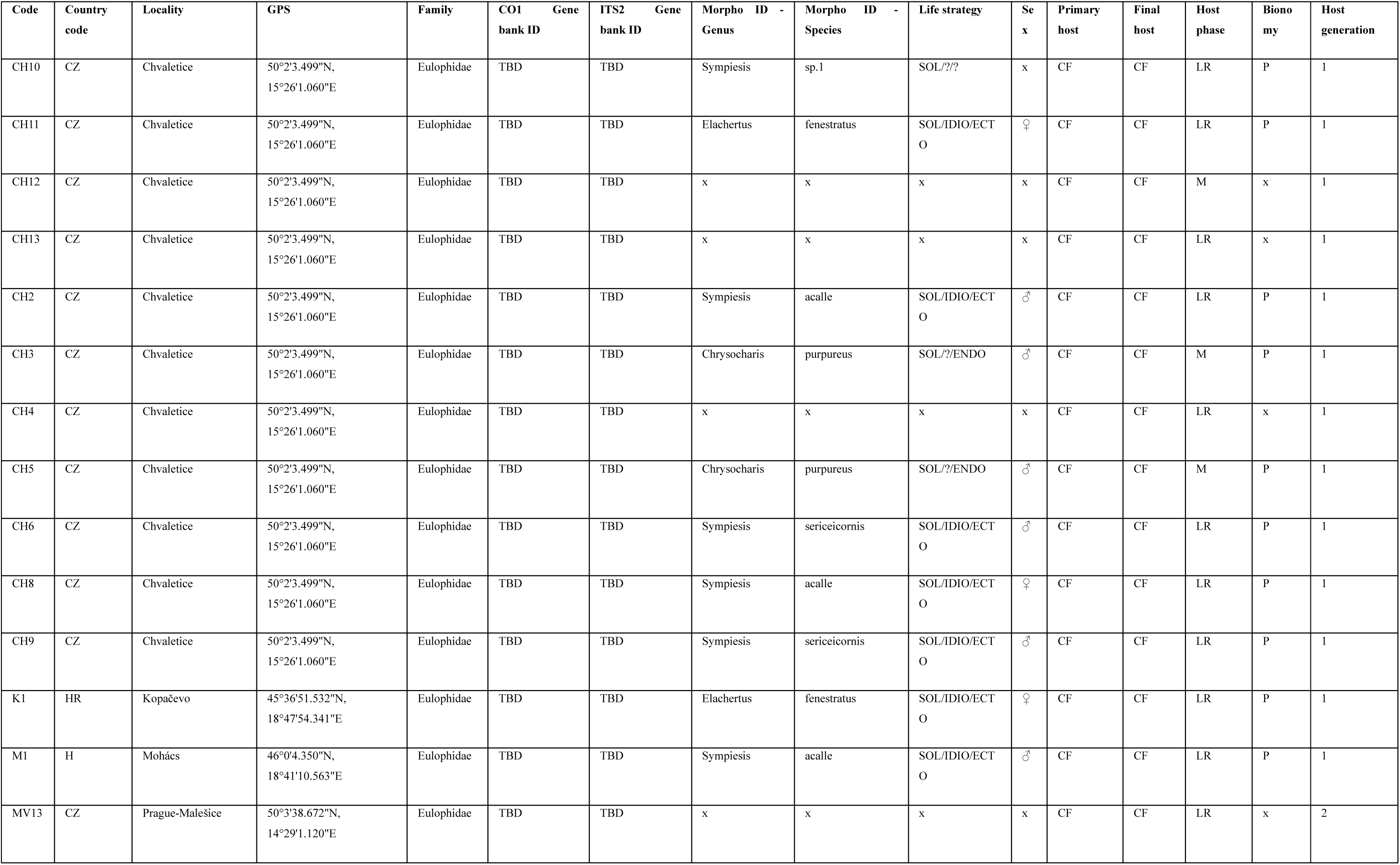
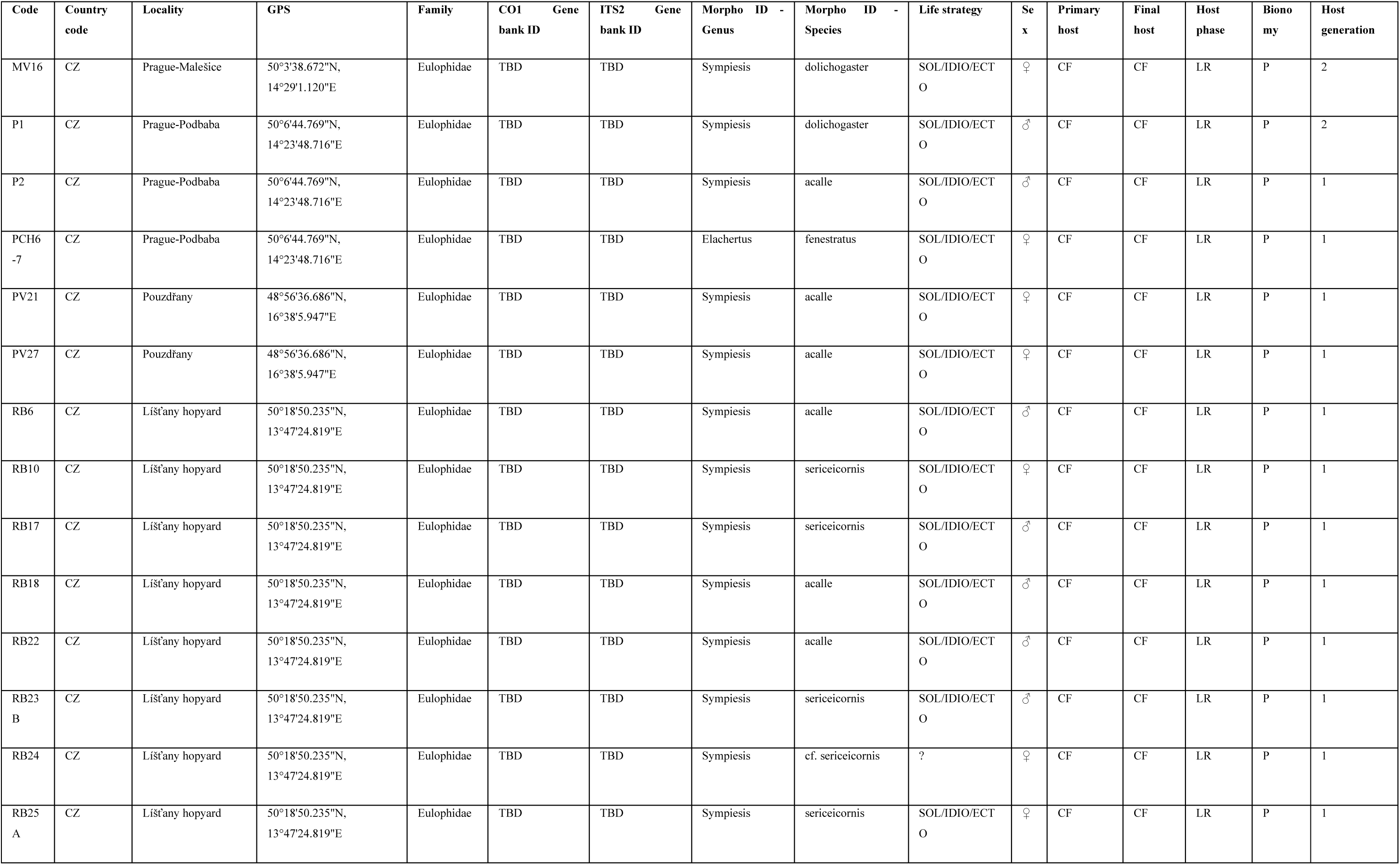
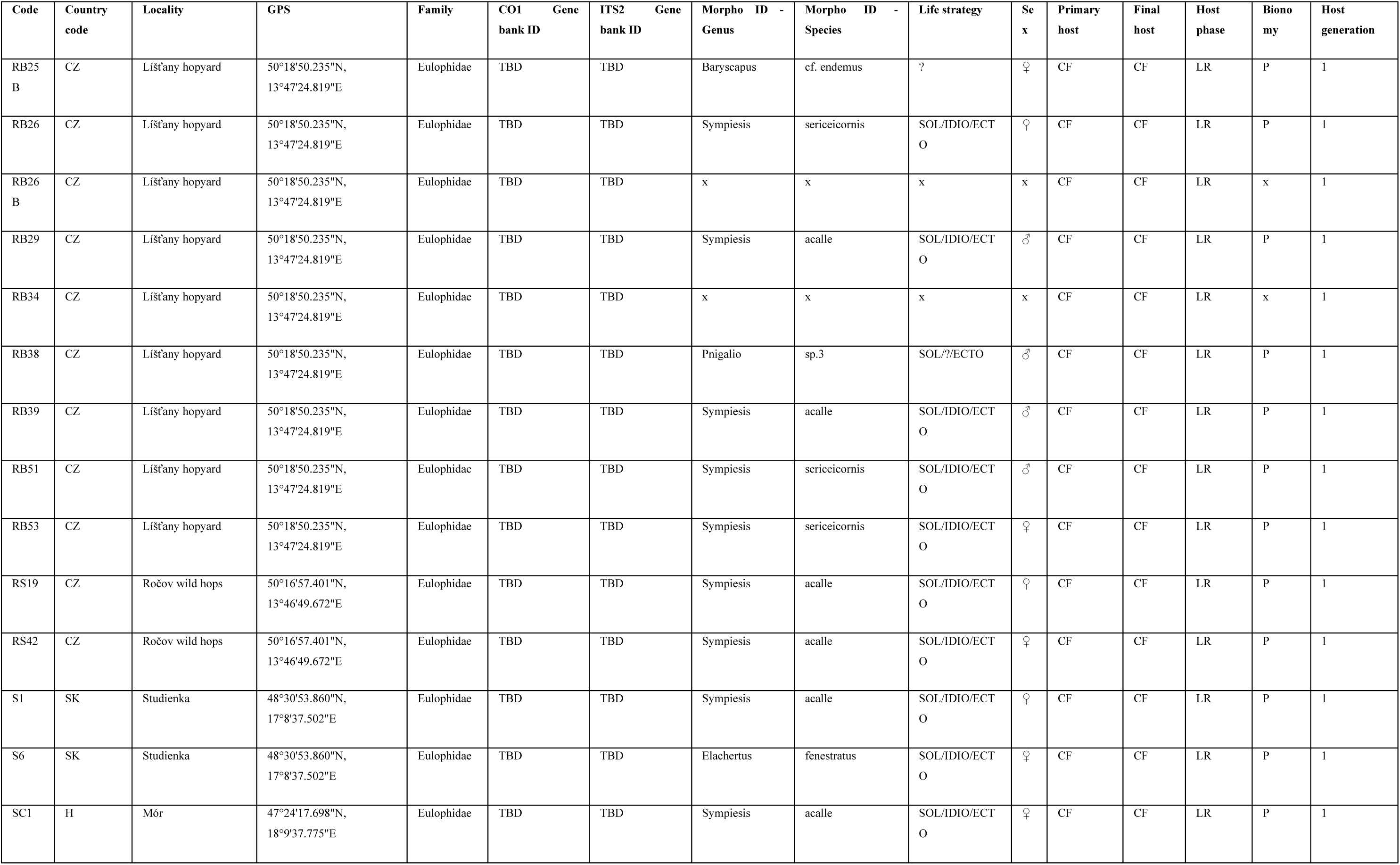
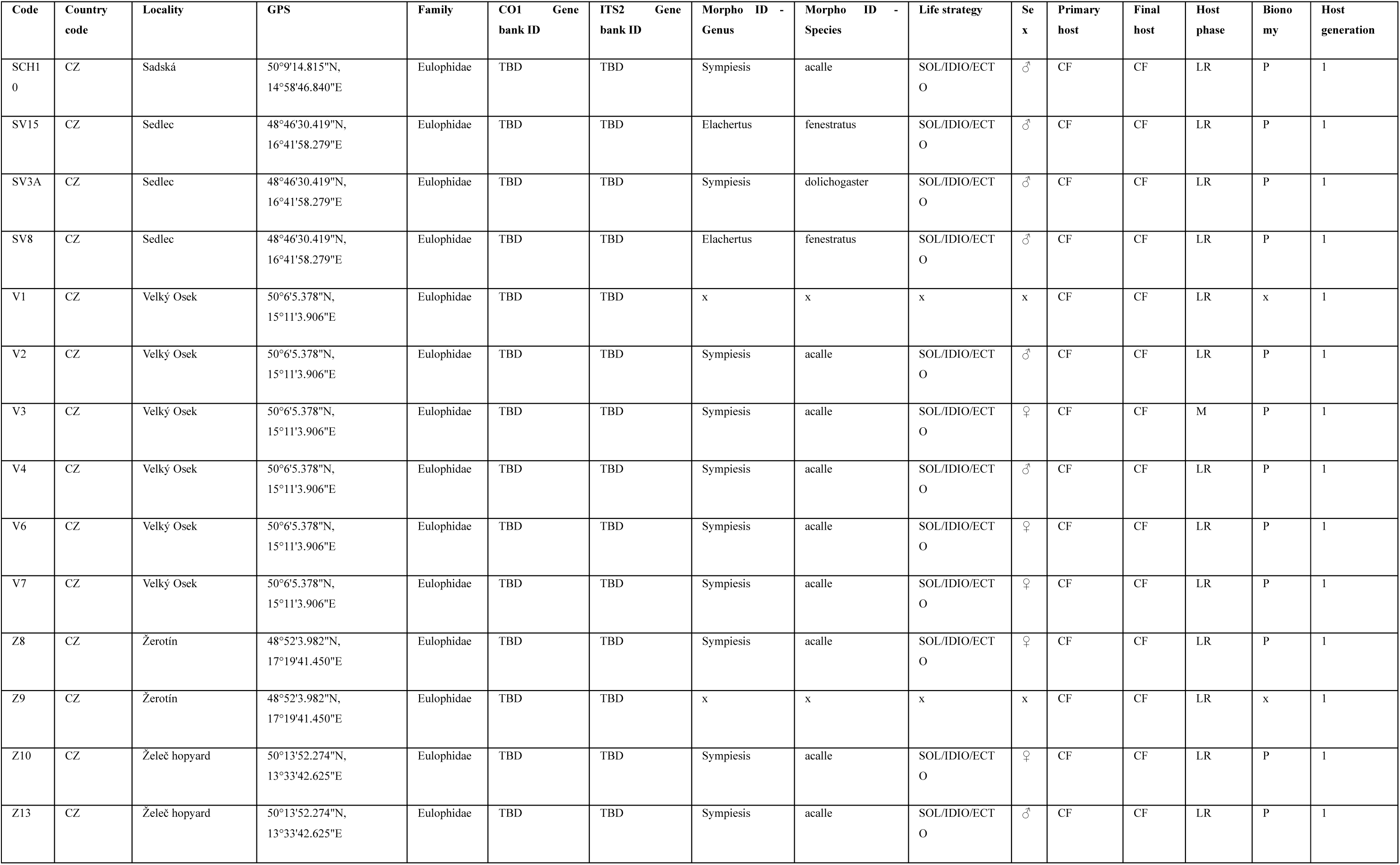
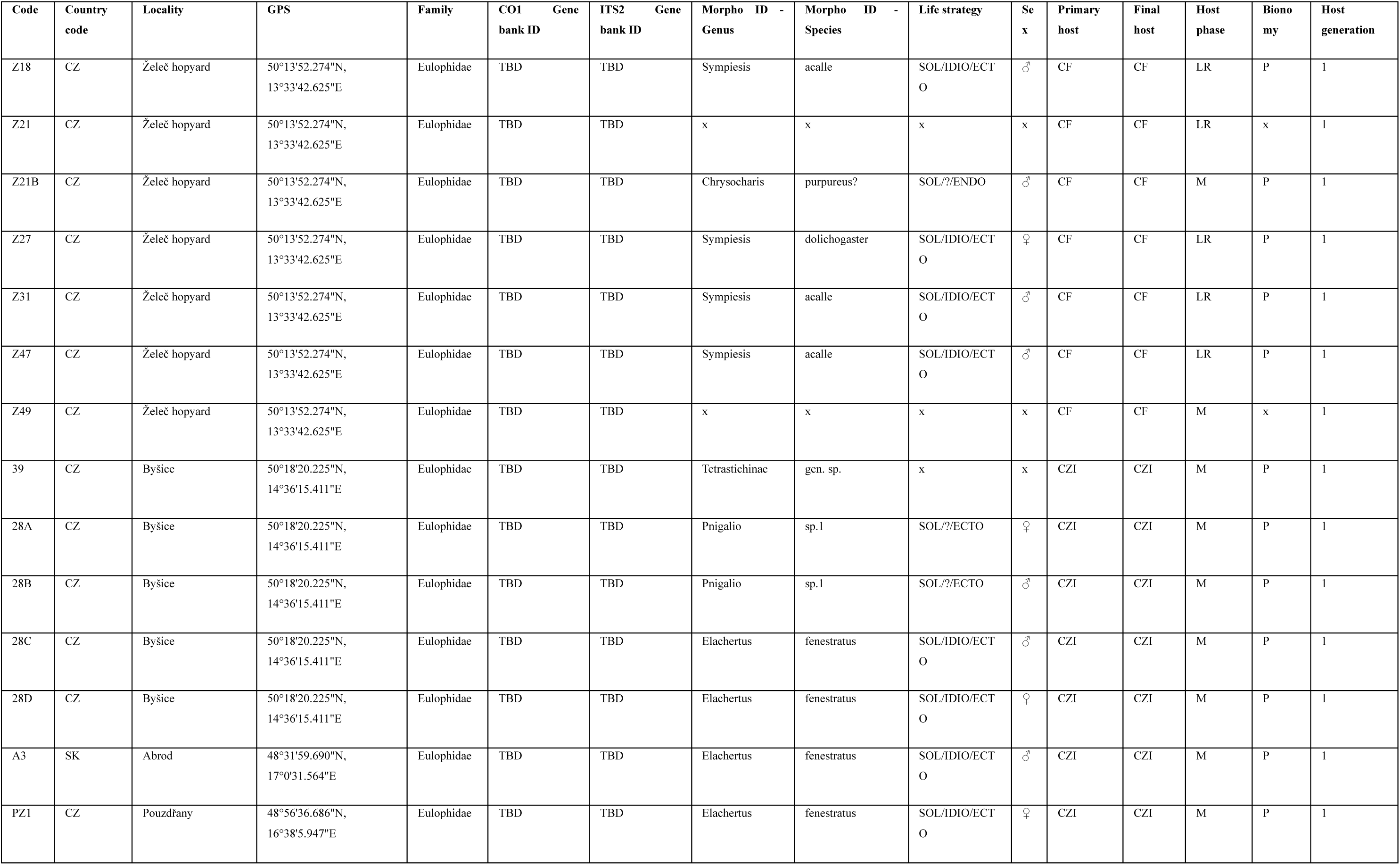
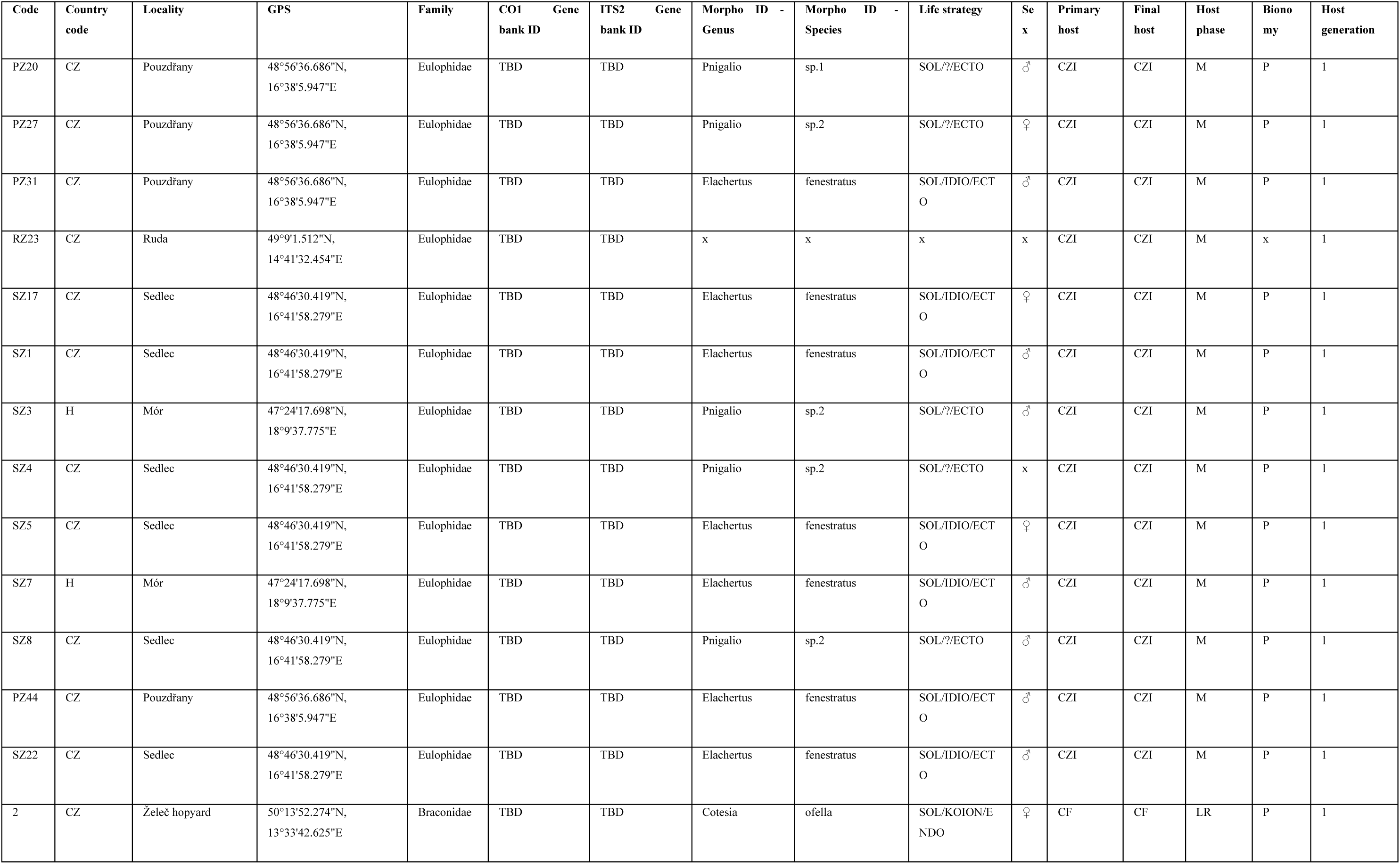
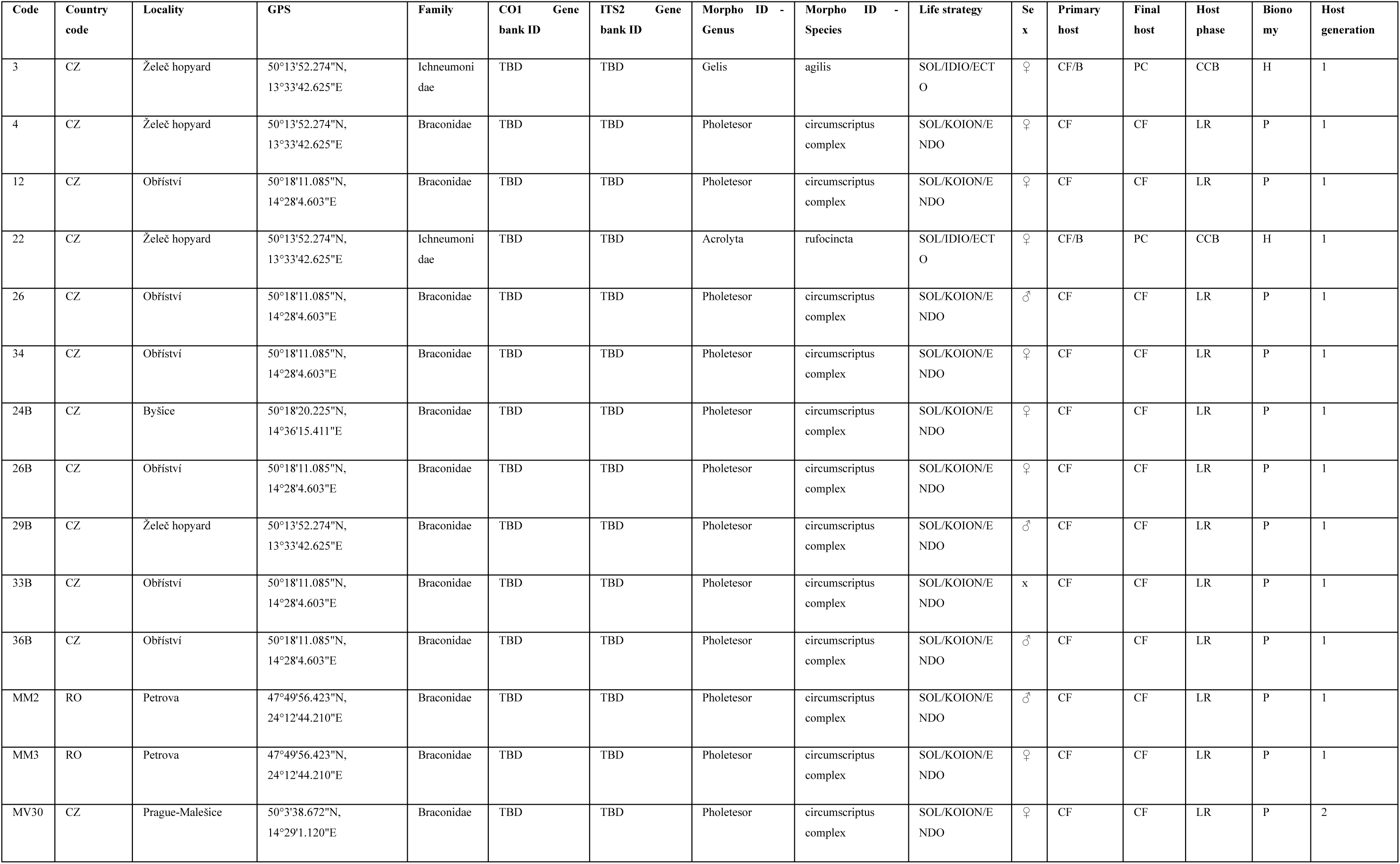
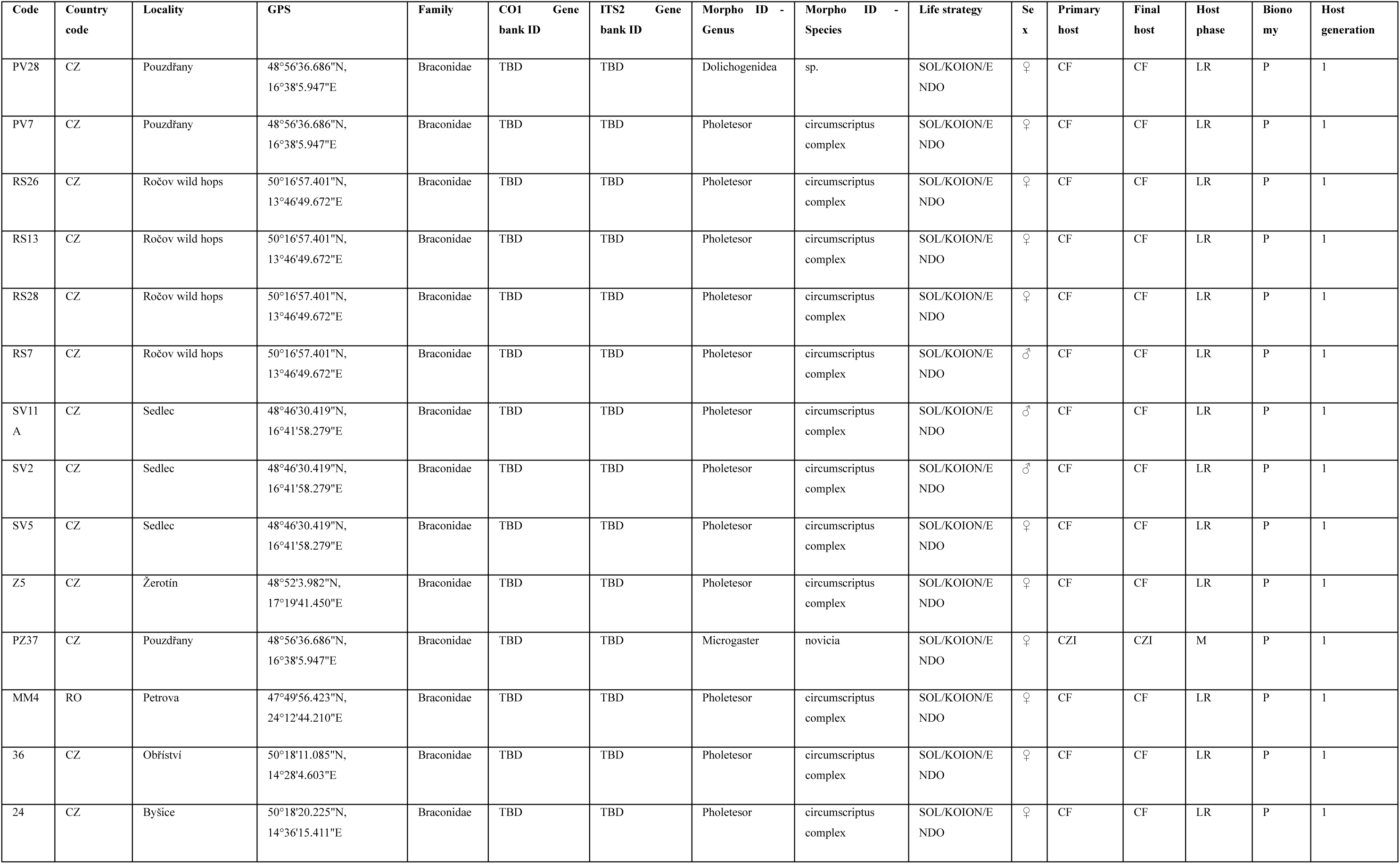
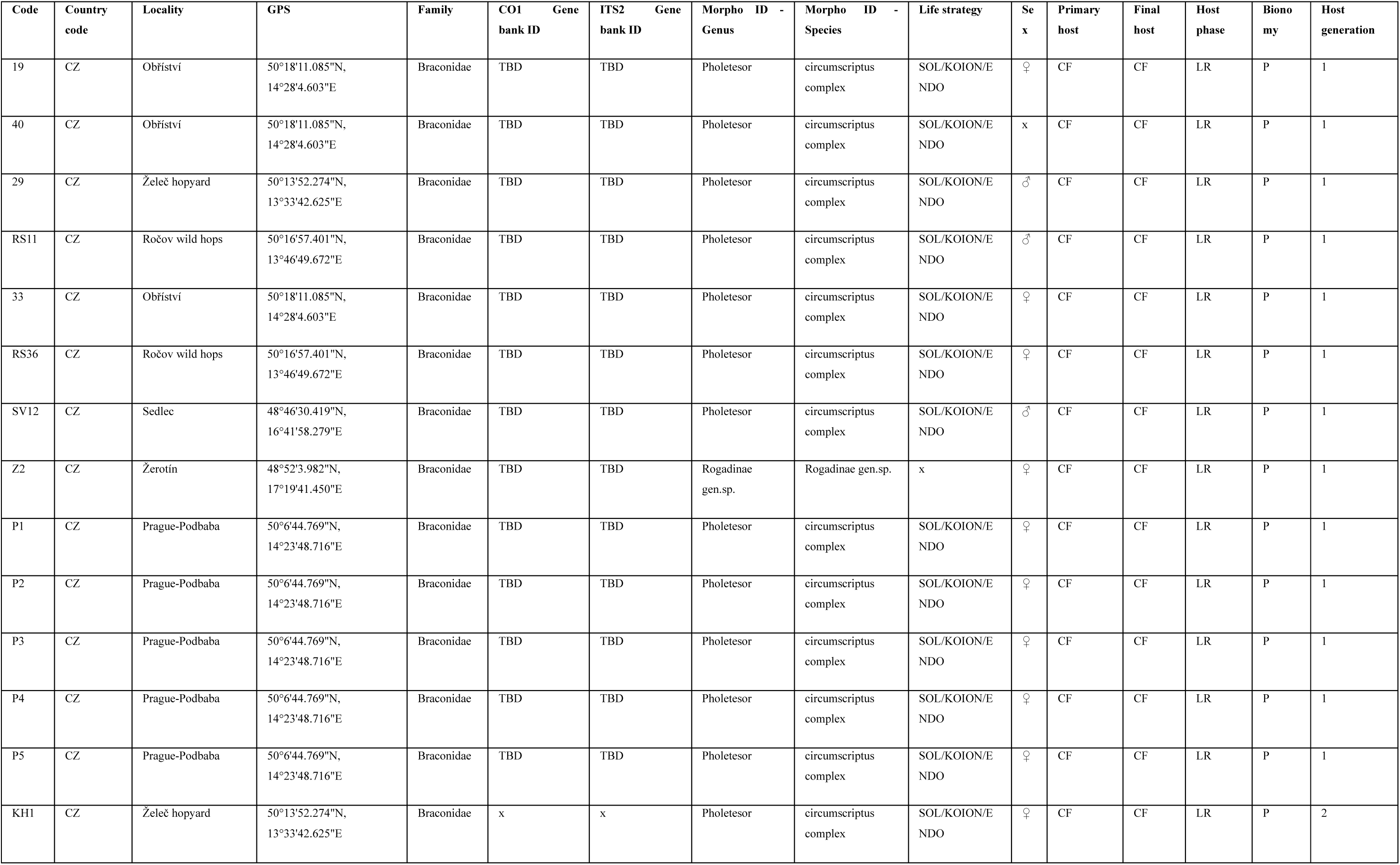
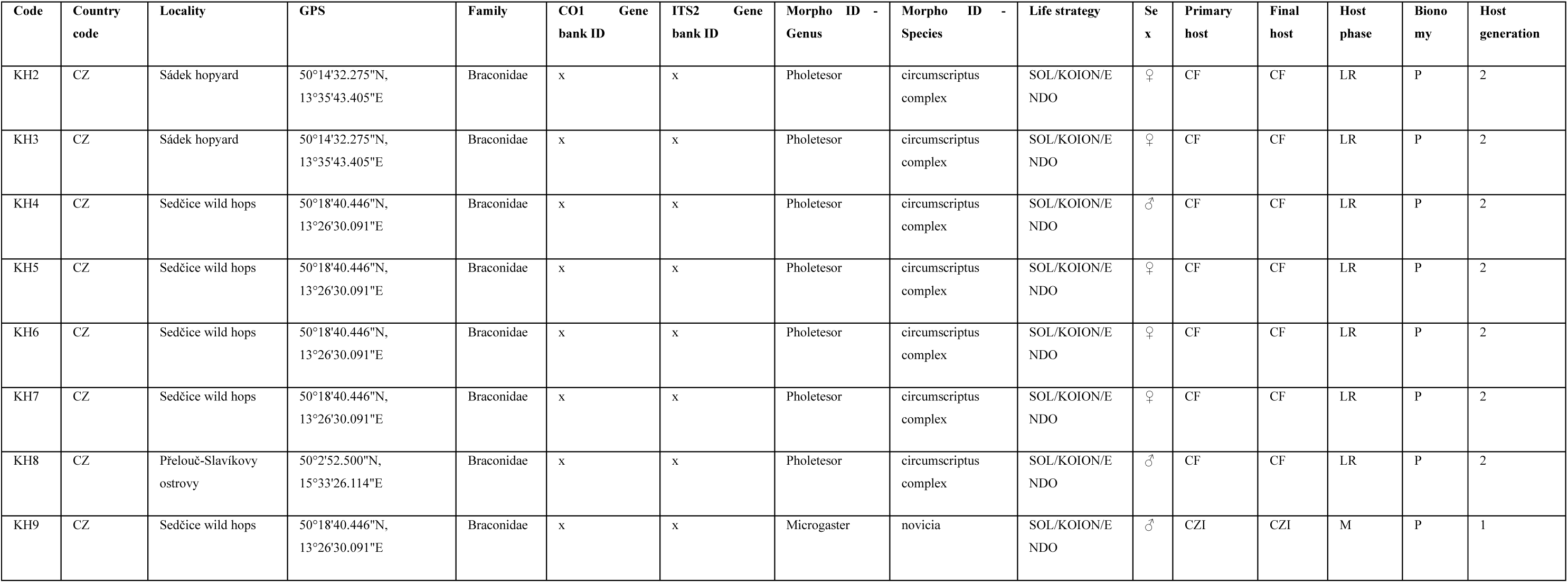
Observed parasitoids of *Caloptilia fidella* and *Cosmopterix zieglerella* recorded in this study. This table presents all parasitoid species recorded in our study, based solely on our observations of *Caloptilia fidella* (CF) and *Cosmopterix zieglerella* (CZI). The table includes information on their life strategy, host association, and developmental characteristics. Abbreviations used in the table: **M** = mine, **LR** = leaf roll, **CCF** = cocoon of *C. fidella*, **CCB** = cocoon of the braconid *P. circumscriptus*, **SOL** = solitary, **IDIO** = idiobiont, **ECTO** = ectoparasitoid, **ENDO** = endoparasitoid, **KOION** = koinobiont, **H** = hyperparasitoid, **P** = primary parasitoid, **P/H** = primary or secondary parasitoid, **PC** = *Pholetesor circumscriptus*, **?** = uncertain or undetected life strategy in our research. The background color in the table represents the host association: yellow shading corresponds to CF, green shading corresponds to CZI, and white cells indicate hyperparasitoids.

### Genetic analysis

DNA was extracted from specimens preserved in 96% ethanol using the Qiagen DNeasy Blood & Tissue Kit with a modified non-destructive protocol based on Cruaud et al. (2019). This protocol was optimized for high DNA yield from small insect specimens while preserving them for morphological identification. Specimens were incubated with lysis buffer and proteinase K at 56°C for 20 hours under gentle vortexing (250rpm). Post-incubation, lysate was transferred to fresh tubes for DNA purification. Elution was performed in two steps using a pre-warmed elution buffer, yielding a final volume of 100 µL with high DNA concentration. Additionally, for high-throughput processing, some samples were extracted using the Xiril AG X100 Automatic Workstation according to Haas et al. (2021) excluding the semi-destructive step as we used the whole specimens. Target gene fragments included CO1 (mitochondrial DNA): LCO1490 (5′-GGT CAA ATC ATA AAG ATA TTG −3′) and HCO2198 (5′-TAA ACT TCA GGG TGA CCA AAA AAT CA −3′) (Hebert et al., 2003) and ITS2 (nuclear ribosomal DNA): ITS2-F (5′-TGT GAA CTG CAG GAC ACA TG −3′) and ITS2-R (5′-AAT GCT TAA ATT TAG GGG GTA −3′) (Campbell et al., 1994). PCR reactions were prepared in 25 µL volumes containing FastGene Optima HotStart Ready Mix (12.5 µL), primers (1 µL), PCR clean H2O (8.5 µL) and template DNA (2 µL). Thermal cycling conditions for CO1 included an initial denaturation at 94°C for 2 minutes, followed by a two-step protocol with annealing temperatures of 45°C for 1 minute (for 5 cycles) and 50°C for 1 minute (for 35 cycles), extension at 72°C for 1.5 minutes for every cycle and for ITS2 amplification followed a similar conditions but used a single annealing temperature of 53°C for 45 seconds (33 cycles). Successful PCR products were purified using ExoSAP-IT™ (1.5µL of PCR clean H2O, 1.5µL of ExoSAP-IT, 1.5µL of PCR product; cycling conditions - 37°C for 15 minutes and 80°C for 15 minutes) and sequenced bidirectionally using Sanger sequencing (Eurofins Genomics, Germany, Ebersberg).

Sequences were processed in Geneious Prime 2023.2.1 (https://www.geneious.com/). Initially, sequences were categorized by taxonomy (Chalcidoidea or Ichneumonoidea), host (*C. fidella* or *C. zieglerella*), and gene (CO1 or ITS2). Forward and reverse of every sequence were assembled, and BLAST was used to determine the taxonomic identity of individuals. Consensus sequences were generated after successful assembly. Alignments were performed in Geneious using the MAFFT plugin with the E-INS-i strategy for ITS2 and L-INS-i strategy for CO1 (Katoh & Standley, 2013), followed by manual verification and trimming to a uniform length. For CO1 sequences, translations to amino acids were checked for stop codons using Geneious for pseudogenes or misalignments. A concatenated alignment of both genes and hosts was created for Chalcidoidea, while for Ichneumonoidea, only the CO1 alignment was used due to the lack of high-quality ITS2 sequences. BLAST was used to verify the taxonomic identity of individuals.

Phylogenetic trees were constructed using the maximum likelihood method in RAxML-HPC2 on XSEDE (8.2.12; Stamatakis, 2006) via the CIPRES server (Miller et al., 2010). The GRTCAT model was used with 1,000 bootstrap replicates (Stamatakis, 2006). Bootstrap percentages (BP) ≥ 70% were considered as strong nodal support. Resulting trees for the concatenated Chalcidoidea dataset and the CO1 alignment for Ichneumonoidea were visualized in FigTree v1.4.4 (Rambaut, 2009). Bayesian analysis was performed in MrBayes v3.2.7 (Ronquist et al., 2012), with input files in NEXUS format set two independent runs with 1,000,000 generations each, saving every 1,000th tree. Parameter files were inspected in Tracer (Rambaut et al., 2018) and the first 25% of trees were discarded as burn-in. Posterior probabilities (PP) ≥ 0.95 were considered as strong support, PP < 0.90 as weak. Final trees were visualized in FigTree, with graphical edits in iTOL (https://itol.embl.de/; Letunic & Bork, 2021).

For species delimitation both gene fragments were used to define species-level entities (OTUs-operational taxonomic units) using two online-available tools. The first method employed was ASAP (Assemble Species by Automatic Partitioning), as described by Puillandre et al. (2021) and Zhang et al. (2022). This method proposes the delimitation of hypothetical species based on genetic distances calculated between DNA sequences. The default settings in the online ASAP interface (available at https://bioinfo.mnhn.fr/abi/public/asap/) were used for the species delimitation analysis. The second method, bPTP, delineated species-level entities using maximum likelihood and the Bayesian implementation of the Poisson Tree Processes (PTP) model for species delimitation (available online at https://species.h-its.org/; Zhang et al., 2013). For the bPTP analysis, the online interface was configured with model settings of 200,000 Markov chain Monte Carlo (MCMC) generations, thinning of 100, and a burn-in of 0.1. ASAP was run on the CO1 and ITS2 sequence alignments in FASTA format without outgroups, while bPTP was run on the phylogenetic trees obtained from these alignments using the maximum likelihood approach.. Additionally, genetic distances across sequences containing CO1 and ITS2 gene fragments were calculated in MEGA 11 using the Kimura-2-parameter (K2P) model (Tamura et al., 2021). Groups of similar sequences were identified using a 2% barcode threshold (Hebert et al., 2003).

### Morphological Identification

To facilitate morphological identification, after lysis every specimen was washed two times in water (15 minutes each bath) and stored in 80% EtOH. Drying was performed according to a modified protocol (Heraty & Hawks, 1998) using hexamethyldisilazane (HMDS). The procedure involved sequential immersion in 90% and 95% EtOH for 30 minutes each, followed by two washes in 100% EtOH for 15 minutes each. Specimens were then placed in HMDS for two 30-minute intervals. After removing HMDS, specimens were left to dry. Due to the volatile nature of HMDS, the process was conducted under a fume hood with appropriate protective equipment. Once dried, specimens were mounted on cards. Prepared specimens were sorted and identified using a Leica M205C stereomicroscope (Leica Microsystems). Identification was based on available keys for Ichneumonoidea (e.g., Nixon, 1965; Whitfield & Wagner, 1991) and Chalcidoidea (e.g., Nikol’skaya, 1952; Bouček, 1959; Yoshimoto, 1983; Gibson et al., 1997).

### Statistical analysis

Data were analyzed using the freely available statistical program R (version 4.1.1) with the FSA library. A non-parametric Kruskal-Wallis test was conducted to determine whether parasitism rates differed among the mine, leaf roll, and pupae stages, as the data did not meet the assumptions of normal distribution. To further explore these differences, a post-hoc Dunn test with Bonferroni correction was applied. Additionally, the Wilcoxon test was used to compare parasitization rates between the two generations.

## RESULTS

### Delimitation of Parasitoid Species

From a total of 162 collected specimens, 139 were successfully sequenced for either CO1 (93 specimens - 67 individuals of Chalcidoidea, 26 of Ichneumonoidea; fragment length ca 622bp) or ITS2 (106 Chalcidoidea; fragment length ranging from 326 to 618bp). For 57 individuals of Chalcidoidea we sequenced both CO1 and ITS2 fragments (summarized in Table 1). Maximum likelihood (RAxML) and BA analyses yielded almost similar topologies for both single locus and concatenate datasets (see Figure 1, Figure S1). Molecular analyses provided significant insights into the identity of parasitoid species associated with *Caloptilia fidella* and *Cosmopterix zieglerella* and allowed corroboration with our morphological hypothesis or identified the specimens that could not be determined based on morphology. Specifically, specimens of Chalcidoidea with codes PZ3, RZ23, 37, MV13, Z9, CH3, V1, RB34, Z21, RB26, CH1, CH12, CH4, MV27, 9, 10 showing severe body damages after DNA extraction were assigned to particular clades based on genetic information.

**Figure 1.**
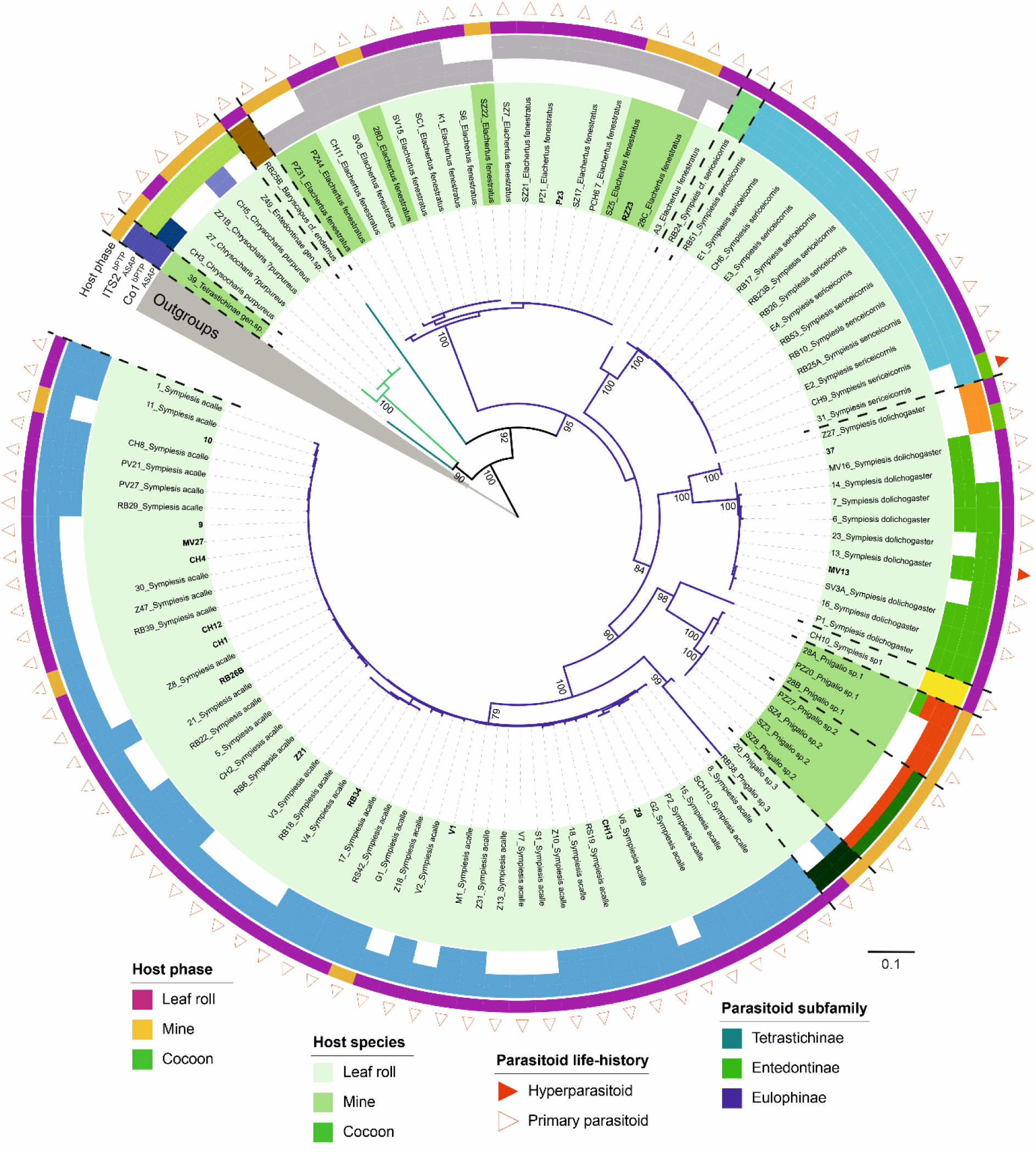
Phylogenetic tree of Chalcidoidea parasitoids based on RAxML analysis of concatenated COI and ITS2 sequences. Bootstrap values >70 are indicated at branch nodes. Branch colors correspond to different families, while the background shading of parasitoid species represents their respective hosts. Additionally, colors distinguish the host life stages parasitized, recorded parasitoid bionomy, and species delimitation based on genetic distances using the ASAP and bPTP methods.

Using morphology, we distinguished 17 species of parasitoids. Within Ichneumonoidea we identified 6 species (2 species of Ichneumonidae and 4 of Braconidae). *Cosmopterix zieglerella* hosted only one species of Braconidae (*Microgaster novicia* Marshall, 1855), while *C. fidella* hosted three species of Braconidae and two species of Ichneumonidae. Chalcidoidea were represented by 11 species (all Eulophidae) with four species of parasitoids of *C. zieglerella* and 8 species of *C. fidella*.

While the morphological analysis identified 17 parasitoid species either on(?) *C. fidella* or *C. zieglerella*, molecular species delimitation did not match completely our morphological identification and occasionally varied in relation to the method used in some parasitoid groups. For Ichneumonoidea, based on CO1, ASAP delimited only 5 putative species, respectively OTUs compared to bPTP or morphology (6 OTU species). For Chalcidoidea, based on COI, ASAP identified a total of 10 OTUs and bPTP 12 OTUs, based on ITS2, ASAP delimited 12 OTUs and bPTP 13 OTUs. Results of delimitation methods have been mapped on the maximum likelihood tree calculated using a concatenated dataset in Chalcidoidea (Figure 1) and CO1 dataset in case of Ichneumonoidea (Figure S2-4).

Both delimitation methods pointed out one probably cryptic species within Chalcidoidea. Specifically, RB24 identified as *Sympiesis sericeicornis* (Nees, 1834) had no morphological differences from other specimens identified as S. *sericeicornis.* The genetic distance between these two clades (RB24 and the rest of *S. sericeicornis*) on CO1 was 8.7% and on ITS2 4.8% and corroborated the hypothesis of the cryptic species. However, we did not decide to describe this specimen (further referred as RB24_*Sympiesis* cf. *sericeicornis* as a new species as it requires more intensive research on more specimens, ideally from other host species/areas.

In a few cases, delimitation methods underestimated or overestimated the number of OTUs. ASAP method considered the two (PV28_*Dolichogenidea* sp. - Braconidae and 28A_*Pnigalio* sp.1 - Eulophidae), morphologically very well distinguishable species, as the same OTUs as their sister clades (i.e., 2_*Cotesia ?ofella* and *Sympiesis dolichogaster* Ashmead, 1888 - in case of CO1, and *Pnigalio* sp.2 - in case of ITS2). However, the genetic distance on CO1 has been discovered to be over 10% (for PV28_*Dolichogenidea* sp.) and 8.7% or 4.2% respectively on ITS2 (for 28A_*Pnigalio* sp.1). Therefore, in this case, we decided to follow the bPTP analysis and recognize into two species. On the other hand, the two specimens of *Chrysocharis purpureus* Bukovskii, 1938 (CH3 and CH5), although morphologically uniform, were delimited as two separate species with both methods using CO1 with quite large genetic distance (8.7%). However, analysis of the ITS2 dataset and concatenated dataset did not show such divergence and also both delimitation methods presented the specimens as one species. The intraspecific distance of other species of parasitoids did not exceed over 2% neither for COI, nor ITS2.

### Parasitoids composition

A total of 162 hymenopteran parasitoids were successfully reared from 924 collected leaves containing hosts (*C. fidella*: 142 individuals of 14 parasitoid species from 774 leaves, overall average parasitization of 18.34%; *C. zieglerella*: 20 individuals of five parasitoid species from 150 leaves, overall average parasitization of 13.33%) on 18 collection sites between 2020-2022 (Table 1). Notably, *Elachertus fenestratus* (Nees, 1834) was the only species associated with both *C. fidella* and *C. zieglerella*. Out of a total of 142 parasitoid individuals associated with *C. fidella*, 122 emerged from leaf rolls, 12 from mines, and 8 from the pupae of the primary host or primary parasitoid. Of those emerging from leaf rolls, three OTUs were identified as koinobionts, six as idiobionts, and the bionomy of five OTUs was not further detailed. For *C. zieglerella*, all 20 parasitoids emerged from the host’s mine. Among these OTUs, one was identified as solitary idiobiont ectoparasitoid, three as possible idiobiont ectoparasitoids and one as solitary koinobiont endoparasitoids (Figure 2).

**Figure 2.**
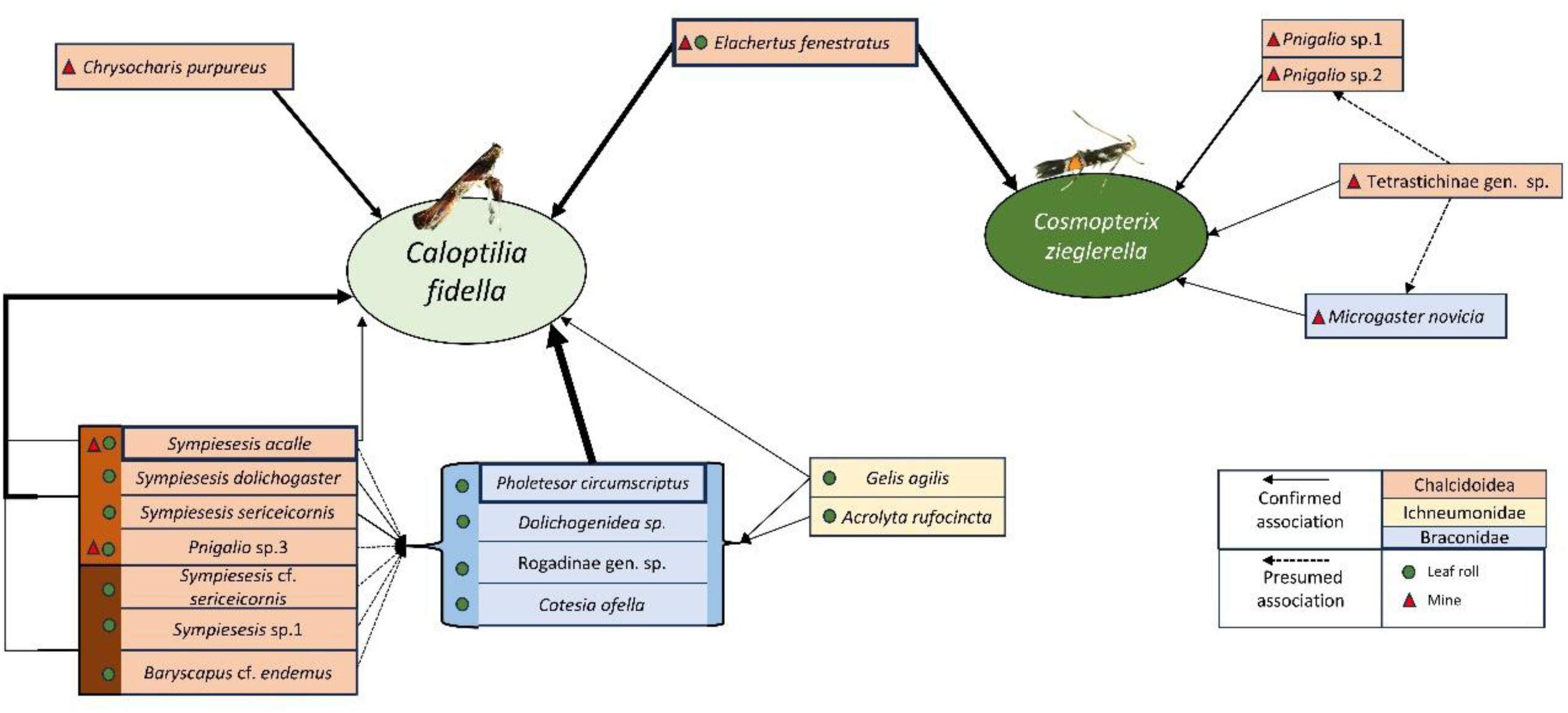
Overview and associations of observed parasitoids in this study with the studied hosts. The thickness of the lines between species indicates the frequency of parasitoid-host associations, i.e., thicker lines represent more frequent associations. Colored symbols indicate the developmental stages of the host from which the parasitoids emerged.

**Figure 3.**
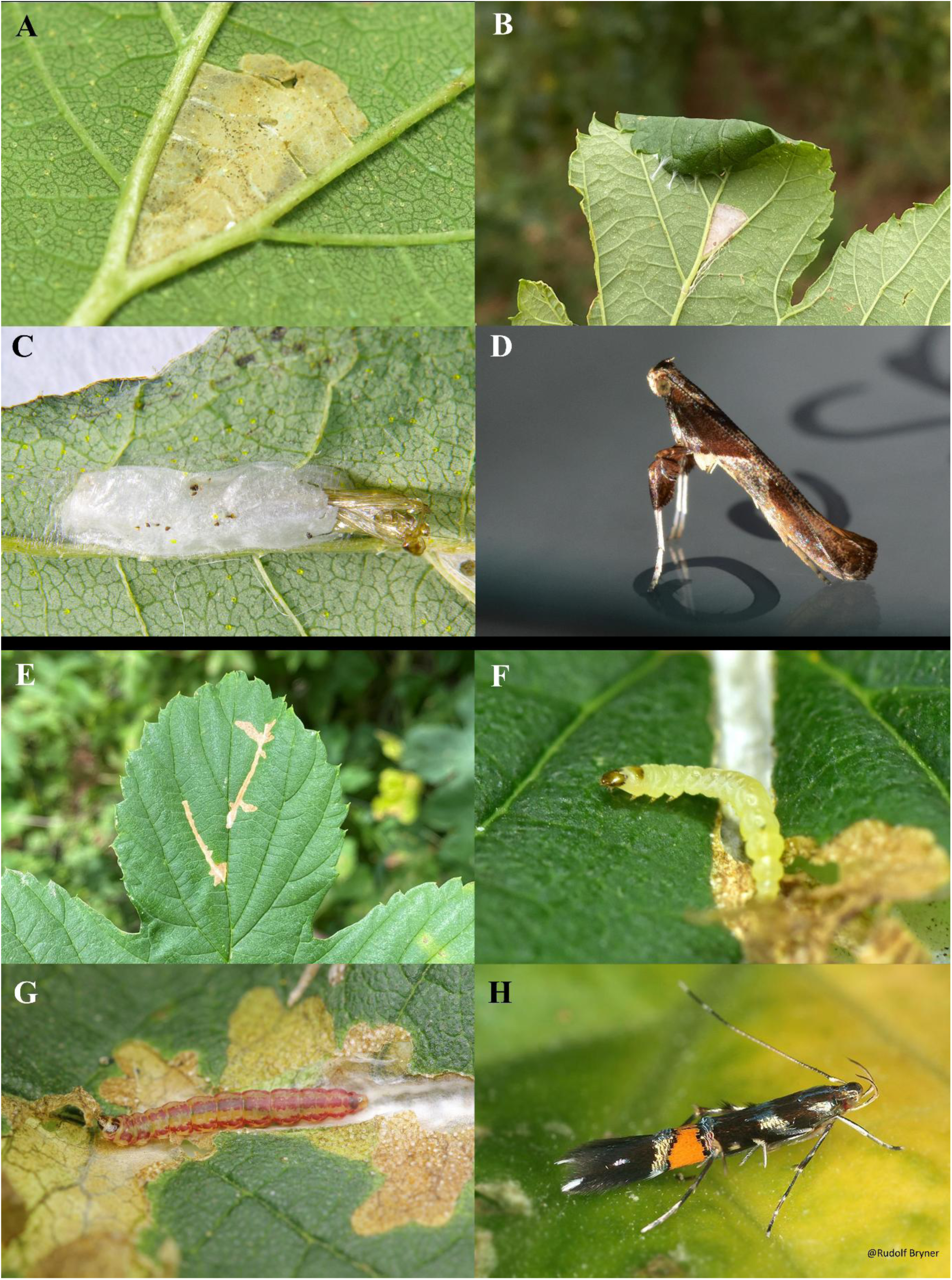
Life cycle of two microlepidoptera living on hops. **A–E.** *Caloptilia fidella*: **A.** Mining larva within a leaf mine located between the veins at the leaf axil. **B.** Leaf mine and a leaf roll with visible silk threads spun by the larva. **C.** Silvery silk cocoon with remnants of the larval pupa on the underside of the leaf. **E–H.** *Cosmopterix zieglerella*: **E.** Characteristic leaf mine on hop leaves, which later develops into a broader, flattened shape. **F.** Young larva. **G.** Final instar larva before pupation. **H.** Adult moth. (Photo credits: T. Hovorka, K. Holý, and image **H** by Rudolf Bryner)

### Parasitoids associated with Caloptilia fidella

Braconidae were represented by two parasitoid species. *Pholetesor circumscriptus* (Nees, 1834) emerged as the predominant primary parasitoid, exhibiting koinobiont endoparasitism of leaf-mining larvae and early exophagous stages within a leaf roll (Figure 4A-B). A total of 39 individuals were reared, with a sex ratio favoring females (27 females to 12 males), highlighting the central role of this species in the community. Another braconid parasitized leaf roll, tentatively identified as *Cotesia ?ofella* exhibiting koinobiont endoparasitism, was represented by a single female.

**Figure 4.**
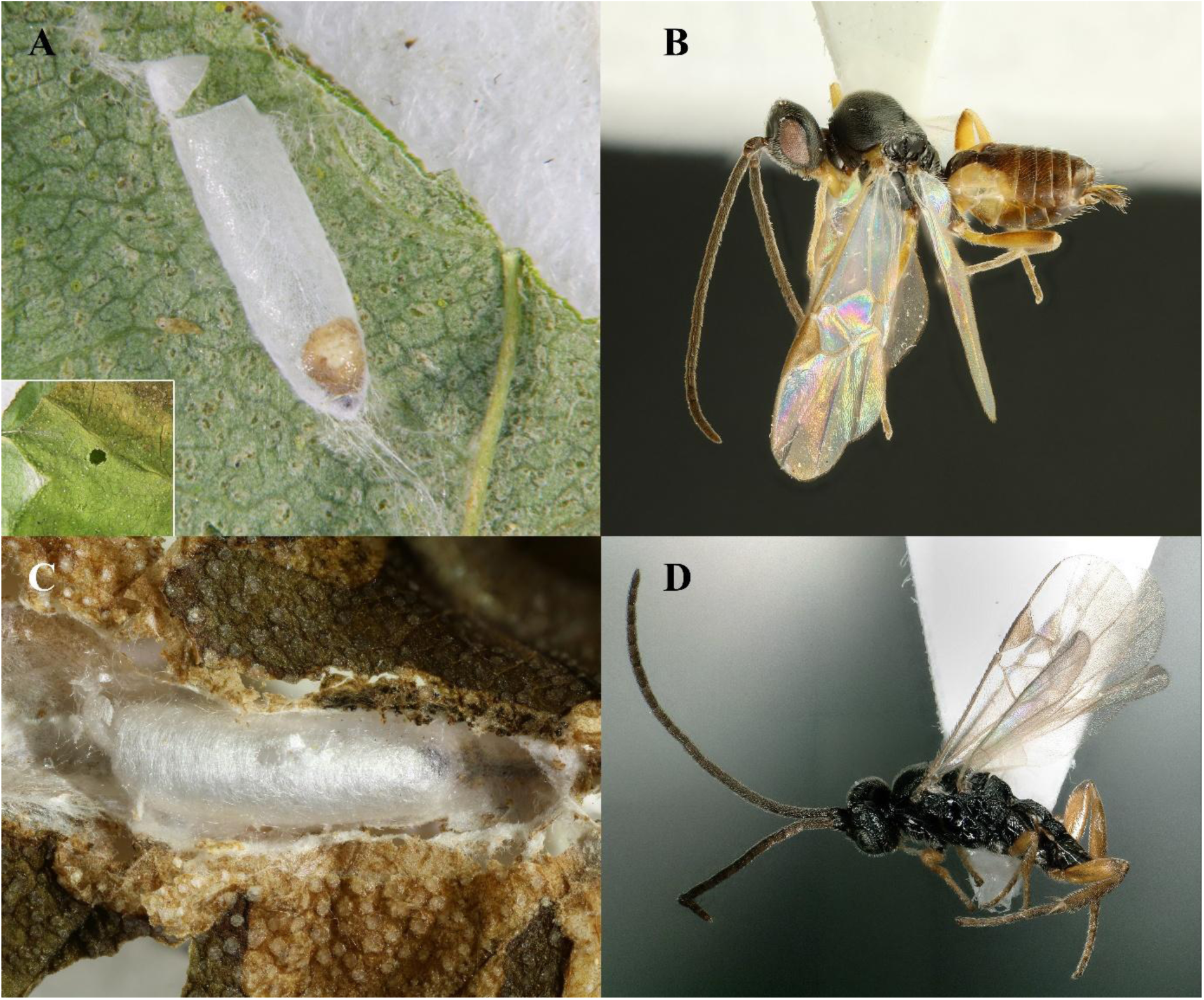
Parasitoids of the subfamily Microgastrinae reared from hosts and identified to species. A. Cocoon with polar threads and an opened top, typical for the parasitoid *Pholetesor circumscriptus* (visible in the bottom left corner is an emergence hole chewed by the parasitoid in the leaf roll of the host *Caloptilia fidella*). **B.** Lateral habitus of *P. circumscriptus* (female) reared from *C. fidella*. **C.** Characteristic silk cocoon of the parasitoid *Microgaster novicia* located inside the mine of the host *Cosmopterix zieglerella*. **D.** Lateral habitus of *M. novicia* (male) reared from the host *C. zieglerella*. (Photo: T. Hovorka)

Within the Ichneumonidae, *Gelis agilis* (Fabricius, 1775) and *Acrolyta rufocincta* (Gravenhorst, 1829) were identified as hyperparasitoids (Figure 5A-B). These species specifically targeted the cocoons of *P. circumscriptus*, contributing to the intricate parasitization dynamics. Notably, *G. agilis* exhibiting solitary idiobiont ectoparasitoidism, emerged from *P. circumscriptus* cocoons, further emphasizing its role in secondary parasitism.

**Figure 5.**
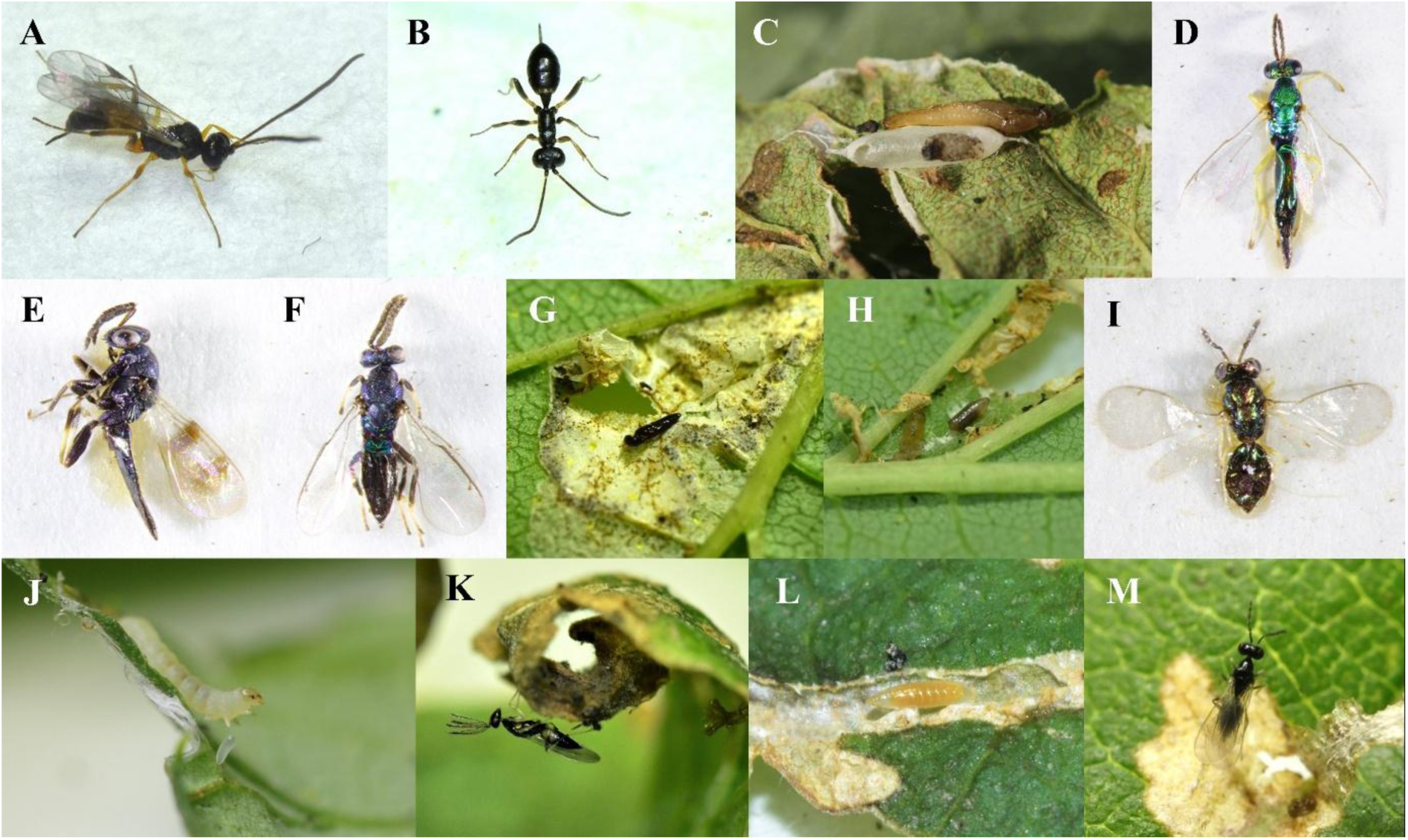
Parasitoids emerged from *Caloptilia fidella* (CF), *Cosmopterix zieglerella* (CZI) and *Pholetesor circumscriptus* (PC). **A.** Acrolyta rufocincta (PC), **B.** Gelis agilis (PC), **C., D.** Sympiesis dolichogaster (PC), **E.** S. acalle (CF), **F.** S. sericeicornis (CF), **G., H., I.** Pupa, larva and adult of *Chrysocharis purpureus* (CF), **J.** Egg near paralized caterpillar of CF, K. *Pnigalio* sp. (CF, CZI), **L.** larva of *Elachertus fenestratus* in mine of CZI, **M.** E. fenestratus emerged from mine of CZI. (Photo: T. Hovorka and K. Holý)

Eulophidae were represented by several species performing with multifaceted ecological roles. *Sympiesis dolichogaster* (Figure 5C-D) and *Sympiesis sericeicornis* (Figure 5F) acted mostly as primary idiobiont ectoparasitoids of leafrolls. However, *S. dolichogaster* also occasionally develops as a hyperparasitoid, attacking prepupae of *P. circumscriptus*. Solitary idiobiont ectoparasitoid *Sympiesis acalle* (Walker, 1848) (Figure 5E) and solitary endoparasitoid *Chrysocharis purpureus* parasitized larvae within mines and leaf rolls (Figure 5G-I), with *S. acalle* ranking as the second most abundant parasitoid after *P. circumscriptus*. Another ectoparasitoid, *Elachertus fenestratus*, primary solitary idiobiont ectoparasitoid, was recorded parasitizing late-instar larvae within leaf rolls.

In addition to the above-mentioned species of parasitoids from the family Eulophidae, additional individuals were reared from leaf rolls containing larvae: *Baryscapus* cf*. endemus* (female, subfamily Tetrastichinae), *Pnigalio* sp.3 (female), and *Sympiesis* sp.1 and *Sympiesis* cf*. sericeicornis* (females, subfamily Eulophinae). One individual of *Pnigalio* sp.3 (male) was reared from a mine (Figure 5K). Precise species-level identification was not possible due to damage to the specimens during either DNA extraction or preparation and CO1 sequences of such OTUs are not presented in GenBank. In the case of the female *Pnigalio* sp.3, a freely laid egg was observed next to a paralyzed larva in leaf roll (Figure 5J). The parasitoid larva fed as a solitary idiobiont ectoparasitoid on the host. The genus *Pnigalio* has not been previously recorded as a parasitoid of *C. fidella* larvae, making this a novel host record. The female *Baryscapus* cf*. endemus* was reared from a leaf roll without detailed observations of its biology. This represents the first known association of the genus *Baryscapus* with the host genus *Caloptilia*. Individuals identified as *Sympiesis* sp.1 and *Sympiesis* cf. *sericeicornis* were reared as primary parasitoids of *C. fidella* larvae within leaf rolls. Genetically, they differ from previously recorded species of the genus *Sympiesis*: *Sympiesis* sp.1 clusters closely with parasitoids of the genus *Pnigalio*, while *Sympiesis* cf. *sericeicornis* is genetically close to species of *S. dolichogaster*.

### Parasitisation of Caloptilia fidella

A total of 774 leaves containing hosts (mines and rolls) were collected, with 534 leaves from the first host generation and 240 from the second host generation. Even if parasitization rates varied significantly across years (Table 2) and developmental stages, no significant difference in parasitization rates was observed between the first and second generations (Wilcoxon test, p-value = 0.4801). However, different larval stages exhibited significant variation in parasitization rates (Kruskal-Wallis test, p-value: 0.0008573). Out of a total of 142 parasitoids reared from *C. fidella*, 122 individuals emerged from leaf rolls (parasitization rates 86%), 12 from mining larvae (8%), and eight from pupae (6%). Larvae within leaf rolls were significantly more parasitized compared to pupae and mining larvae (post-hoc Dunn test, p-values: 0.007864636 and 0.009008146, respectively). Pupation stages exhibited the lowest parasitization (6%), primarily involving morphologically unidentified Eulophidae (in 74%; four of them were damaged during DNA isolation, and no sequence of those was obtained), *Sympiesis dolichogaster* (13%) and *S. sericeicornis* (13%). Mining larvae had the second lowest parasitization rates (8%), primarily caused by *C. purpureus* (in 33%), *Sympiesis* species (in 22%), *Pnigalio* sp.3 (in 11%) and three morphologically unidentified OTUs (in 33%; destroyed during DNA isolation). The first and second exophagous larval stages were the most susceptible, with parasitization rates reaching 86%, largely attributable to *P. circumscriptus* (in 32%) and *Sympiesis acalle* (in 31%).

**Table 2.**
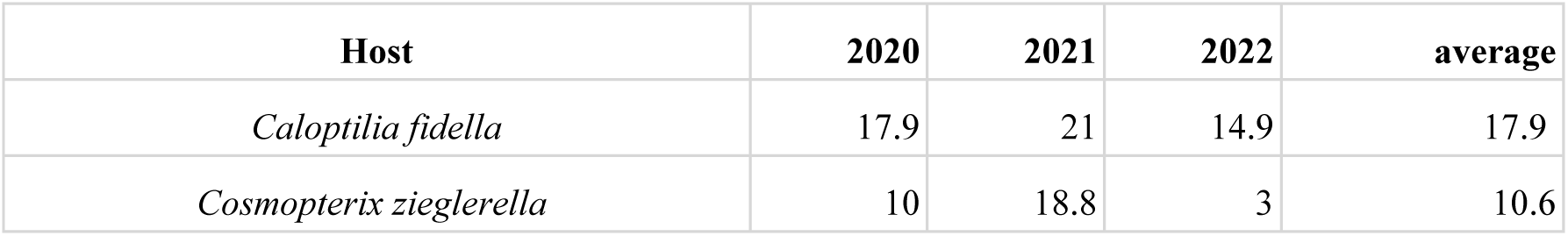
Total and average parasitism rates of individual hosts recorded in this study.

| Host | 2020 | 2021 | 2022 | average |
| --- | --- | --- | --- | --- |
| *Caloptilia fidella* | 17.9 | 21 | 14.9 | 17.9 |
| *Cosmopterix zieglerella* | 10 | 18.8 | 3 | 10.6 |

### Parasitoids associated with Cosmopterix zieglerella

Braconidae were represented by *Microgaster novicia*, which was observed as a solitary, koinobiont endoparasitoid of the mining larvae of *C. zieglerella* (Figure 4C-D). Only a single female of this species was reared from the samples. The larva of *M. novicia* emerged from a final (purple) instar larva of *C. zieglerella*, which exhibited no outward signs of parasitism until shortly before pupation. The parasitoid cocoon, formed within the host’s mine, was distinctly different from that of *P. circumscriptus* in its absence of polar filaments (Figure 4).

Within Eulophidae, *Elachertus fenestratus* emerged as the most abundant parasitoid of *C. zieglerella* (Figure 5L-M). This primary, solitary idiobiont ectoparasitoid was observed in 11 individuals (four females and seven males), accounting for a significant proportion of parasitation events (52%). Detailed observations revealed that females of *E. fenestratus* initially paralyzed their host larvae and subsequently consumed them within the mines. In several cases, larval parasitoids were seen actively moving within the mine to avoid light exposure. Other eulophid parasitoids included individuals of *Pnigalio* sp.1 and *Pnigalio* sp.2 and an unidentifiable individual of the subfamily Tetrastichinae gen. sp.1. All specimens of this species were poorly preserved, limiting precise identification. In at least one case, *Pnigalio* sp. was observed laying eggs near a paralyzed host larva, suggesting a potential role as a hyperparasitoid. The individual was damaged during DNA isolation, gene amplification failed for both fragments, and precise morphological identification was not possible. Therefore, this individual, along with others of the genus *Pnigalio*, is not assigned to a particular species.

### Parasitization of Cosmopterix zieglerella

A total of 150 leaves containing hosts were collected. Of this total number of leaves, 20 were parasitised. This host has only one generation per year and does not form leaf rolls during its development. Therefore, all developments took place in the mine. For this reason, tests for differential parasitisation between generations and life stages were not performed for this host species. The parasitization rates for individual years are in Table 2. A total of 20 parasitoids were reared across the three years, with solitary emergence observed in all cases. Mining larvae were predominantly parasitized by *E. fenestratus* (in 52%), which displayed an ectoparasitic strategy. No parasitoid has been reared from host pupae.

### Hyperparasitism and Novel Findings

For *C. fidella*, hyperparasitism was a significant factor shaping the parasitoid community. *P. circumscriptus* cocoons were frequently attacked by chalcidoid and ichneumonid hyperparasitoids, with *S. dolichogaster* exhibiting dual roles as a primary parasitoid of *C. fidella* and hyperparasitoid of *P. circumscriptus*. For *C. zieglerella*, no direct hyperparasitism was observed. However, the role of *Pnigalio* sp. as potential hyperparasitoids warrants further investigation.

This study uncovered several novel host-parasitoid associations and newly identified *S. acalle*, *S. sericeicornis*, and *E. fenestratus* as parasitoids with an idiobiont life strategy. For *C. fidella*, these included the first records as a host for *P. circumscriptus*, *S. dolichogaster* and *S. acalle*. All parasitoids of *C. zieglerella* were recorded for the first time and therefore represent completely novel findings.

## DISCUSSION

In previous studies, the families Eulophidae (Chalcidoidea) and Braconidae (Ichneumonoidea) have been consistently reported as one of the most diverse parasitoid wasp groups associated with leaf mining insects (Salvo et al., 2011). Our study confirms this. Morphological identification of these dark taxa is challenging, and many lineages might represent cryptic species (Schmidt et al., 2015, 2020) that are not possible to distinguish based on morphology only. However, to reveal the complexity of trophic interaction between parasitoids and their hosts requires the most precise species identification for further understanding of life histories and evolutionary consequences between parasitoids and their hosts or host plants. Therefore, besides morphology, we also applied barcoding and species delimitation. Such an integrative approach has been widely used to successfully decipher the identity of various groups of insects without excluding the parasitoids (Zhang et al., 2013, 2022; Sheikh et al., 2022; Budrys et al., 2023).

The two molecular markers used in this study, ITS2 and CO1, provided complementary data for species delimitation. However, ITS2 was particularly effective in Chalcidoidea, offering greater amplification success than CO1. ITS2 is increasingly recognized as a valuable marker for distinguishing closely related parasitoid species, as demonstrated by studies on various parasitoid families, including Microgastrinae (Fagan-Jeffries et al., 2018) and Eulophidae (Perry & Heraty, 2019). The results of this study further confirm the utility of ITS2 for resolving taxonomic challenges, particularly within morphologically similar taxa.

In our study, both methods (ASAP and bPTP) aligned relatively well with morphological identity of parasitoids with a few exceptions. While in some cases, ASAP underestimated, the number of OTUs identified using morphology (i.e., 28A_*Pnigalio* sp.1 – on CO1 and ITS2, and 20_*Pnigalio* sp.3 – on CO1 only), bPTP proved to be more sensitive. It identified a higher number of OTUs than we were able to distinguish based on morphological differences (i.e., RB24_*Sympiesis* cf. *sericeicornis* – on CO1 and ITS2, and 37 + Z27_*Sympiesis dolichogaster* – on ITS2). Underestimation using ASAP might be caused by lack of sufficient number of samples in particular clades (Ahrens et al., 2016; Eberle et al., 2020; Vogel et al., 2024). On the other hand, overestimating of OTUs compared to morphology using bPTP has been corroborated by genetic distances on particular gene fragments (RB24_*Sympiesis* cf. *sericeicornis* to the rest of S. *sericeicornis* - 8,8% on CO1 and 4,8% on ITS2, 37 + Z27_*Sympiesis dolichogaster* to the rest of *S. dolichogaster* - 11,5% on ITS2) as the use intraspecific genetic distance on CO1 is set to 2% (Hebert et al., 2003). This might point out to cryptic species within particular clades that we, using current taxon sampling and morphology, are not able to resolve.

The performance of bPTP in this study contrasts with findings from other research, such as Sheikh et al. (2022) and Zhang et al. (2022), which also applied bPTP and ASAP to identify cryptic species in parasitoid complexes. Both studies reported that bPTP tended to overestimate species richness, likely due to its higher sensitivity to subtle genetic variation. For example, Zhang et al. (2022), in their investigation of cryptic species within the genus *Sycophila* Walker, 1871 (Chalcidoidea), and Sheikh et al. (2022), in their study of the *Ormyrus labotus* Walker, 1843 complex (Chalcidoidea), observed discrepancies between the methods, particularly in populations with minor genetic differences or distinct geographic distributions. Both studies relied on the more conservative results from ASAP, suggesting that bPTP might sometimes reflect population-level genetic structuring rather than species-level divergence. The more accurate performance of bPTP in this study can be attributed to the larger genetic distances observed among the parasitoid species associated with *C. fidella* and *C. zieglerella*. These larger intraspecific distances likely reduced the chances of overestimation by bPTP. Moreover, the calculated intraspecific and interspecific genetic distances supported species delimitation via bPTP, as evidenced by phylogenetic analyses.

Despite these successes, some limitations were noted. The CO1 dataset was incomplete, lacking sequences for taxa such as *Pnigalio* sp. 2 and *Pnigalio* sp. 3. This discrepancy limited direct comparisons between ITS2 and CO1. For future studies, obtaining datasets with equal taxonomic representation across both genes would allow for a more robust evaluation of the performance of these molecular markers.

The results of this study highlight the importance of integrating molecular methods with a traditional morphological approach. While molecular techniques are powerful, they can sometimes yield overestimated or ambiguous results if used in isolation. Sharkey et al. (2021) reported cases where molecular methods led to misclassification due to insufficient integration with morphological and ecological evidence. Therefore, an integrative approach combining molecular, morphological, and ecological data is essential for accurate species delimitation (Ahrens et al., 2021; Wang et al., 2024). In our study, molecular methods were used as a complementary tool to confirm morphological identification. Additionally, they proved crucial in cases where certain individuals exhibited slightly different morphological traits, helping us determine that they most likely represent cryptic species that cannot be distinguished morphologically. This was the case, for instance, with *Ch. purpureus* or an individual showing minor differences, designated as RB24_*Sympiesis* cf. *sericeicornis*. Beyond their role in species identification, molecular methods such as ASAP and bPTP have broader applications in ecological and evolutionary research, aiding in biodiversity assessment and species delimitation (Cruaud et al., 2013; Zhang et al., 2013; Sheikh et al., 2022). As sequencing technologies continue to advance, molecular approaches will become even more instrumental in uncovering cryptic diversity and elucidating parasitoid evolutionary relationships (Kenyon et al., 2015).

### Parasitoid community of Cosmopterix zieglerella and Caloptilia fidella

Even if larvae of both hosts are monophagous on the same host plant (*Humulus lupulus*) (Koster & Sinev, 2003; Laštůvka et al., 2018; Watson et al., 2021), *Elachertus fenestratus* is the only parasitoid species shared between the two host species. Except from families Tortricidae, Gelechiidae and Coleophoridae, *E. fenestratus* has been recently referred to as an ectoparasitoid of leaf mining larvae of *Phyllonorycter issikii* (Kumata) (Lepidoptera: Gracillariidae) from *Tilia platyphyllos* (Kosheleva et al., 2022). In Japan, Sugiura (2011) recorded additional *Elachertus* species (*E. inunctus* (Nees, 1834) and another unidentified species) for *Caloptilia azaleella*. Similarly, Barry et al. (2010) in the USA reported another species of *Elachertus* as a parasitoid of *C. porphyretica*. These findings suggest that despite geographical differences, parasitoids of the genus *Elachertus* are common leaf-miner associates. In general, the genus *Elachertus* is known as primary, often gregarious, parasitoid of various Lepidoptera larvae (Schauff, 1985), but our observation confirmed *E. fenestratus* as strictly primary solitary idiobiont ectoparasitoid on both studied hosts either in mines (CZI) or leaf-rolls (CF). Until now, no parasitoids had been recorded for *Cosmopterix zieglerella*. This study reveals the first parasitoid associations for this species, identifying five distinct parasitoid-host interactions. Beside *Elachertus fenestratus,* the other most common parasitoid species included two species of *Pnigalio* (*Pnigalio* sp.1 and *Pnigalio* sp.2 both solitary ectoparasitoids), and one representative of Braconidae, *Microgaster novicia*, solitary koinobiont endoparasitoid (Fernandez-Triana et al., 2020; Shaw, 2023). Other observations of the parasitoid community of *Cosmopteryx* were made by El-Serwy (2006, 2009) in his studies on *C. pararufella* Riedl, 1976 and *C. salahinella* Chrétien, 1907, where *Pnigalio* spp. were among the dominant parasitoids. Additionally, *Cotesia ruficrus* (Haliday, 1834) was frequently recorded in these studies. Ahmad et al. (2022) described a novel parasitoid species, *Bracon cosmopteryx* Ahmad and Pandey, 2022 from *C. phaeogastra* (Meyrick, 1917). These findings indicate that parasitoid complexes of *Cosmopterix* species remain largely unexplored, with only sparse information available for European species of this genus.

The most frequently observed parasitoids of *Caloptilia fidella* were *Pholetesor circumscriptus* (Braconidae) and three species of the genus *Sympiesis* (*S. acalle, S. dolichogaster* and *S. sericeicornis*) (Eulophidae). While *S. acalle* and *S. sericeicornis* are widely distributed across the whole Holarctis, *S. dolichogaster* has been mentioned apart from Nearctic and western Palearctic also in Oriental and Australian realm.

Our study confirms *P. circumscriptus* as a common parasitoid of *Caloptilia fidella*, aligning with previous findings that species of *Pholetesor* frequently parasitize Gracillariidae (Mason, 1981; Papp, 1983; Whitfield, 2006; Liu et al., 2016). In European habitats, *Pholetesor* spp., particularly those within the *circumscriptus* species group, are among the most frequent parasitoids of gracillariid leafminers (Whitfield, 2006; Ahmad et al., 2020), contributing significantly to the natural regulation of these herbivores. These wasps exhibit a koinobiont endoparasitoid strategy, in which the female oviposits into an active leaf-mining larva that continues to develop until the parasitoid reaches its final instar and ultimately kills the host at or near pupation. This developmental strategy is typical of Microgastrinae and allows *Pholetesor* wasps to exploit their hosts efficiently before emergence (Fernandez-Triana et al., 2020). The larvae of *Pholetesor* construct suspended cocoons within leaf mines or leaf rolls, a behavior that has been observed in *P. circumscriptus* and is thought to provide protection against predation or hyperparasitation (Whitfield & Wagner, 1991; Whitfield, 2006). Some *Pholetesor* species, such as *P. pedias*, have also been explored for biological control against agricultural pests (Whitfield, 2006). Recent molecular studies have revealed substantial cryptic diversity within *Pholetesor*, with several genetically distinct lineages hidden within morphologically similar species (Fernandez-Triana et al., 2020). However, in our study, both molecular delimitation methods confirmed that all analyzed specimens belong to *P. circumscriptus*, with no evidence of cryptic diversity. This suggests that *P. circumscriptus* represents a genetically homogeneous species, at least within the studied population.While *Sympiesis dolichogaster* and *S. sericeicornis* were found in our study predominantly as primary ectoparasitoids mostly of leafrolls, *Sympiesis acalle* acted as solitary ectoparasitoid of larvae within either mines or leaf rolls. All three species have been observed here as idiobionts, however, the only so far published mention about idiobiont lifestyle came from study of *S. sericeicornis* on *Phyllonorycter comparella* (Gracillaridae) in Hungary (Szöcs et al., 2015). *S. scalle* has been referred to as a primary ectoparasitoid of larvae or pupae of insects that live in sheltered situations, such as miners and rollers (Maier & Hansson 2006). The species develop rather as generalist and is known from several species of *Caloptilia* and other genera of Gracillariidae, and additionally from many other lepidoptera families such as Gelenichidae, Tischeridae and Tortricidae (Maier & Hansson, 2006; Bouček & Askew, 1968). The two other species have been reported from more distantly related lepidopteran hosts and are considered as generalists, too. The host spectrum of *S. dolichogaster* either includes concealed larvae of Gelenichidae, Gracillariidae, Tischeridae or exposed larvae of Tortricidae (Bouček & Askew, 1968, UCD Community). *S. sericeicornis* is commonly known as primary parasitoid of plethora species of genera Gracillariidae (e.g., *Calloptilia*, *Lithocolletis*, *Phyllonorycter*) and Tischeridae (Bouček & Askew, 1968; Hansson, 1987).

Besides exhibiting a life style as primary parasitoids, these three generalistic eulophids species have also been observed as facultative hyperparasitoids of various braconids including *Pholetesor circumscriptus* (Maier & Hansson, 2006; Hagley, 1985; Maier, 1988; Bouček & Askew, 1968), which corresponds to the results of our study. *Sympiesis acalle* and *S. sericeicornis* have also been recorded as hyperparasitoids of a few Eulophidae such as *Achrysocharoides* or *Pnigalio* (Bouček & Askew, 1968, Askew & Shaw, 1979)

Our findings align with the study by Sugiura (2011), who examined parasitoid assemblages of two other *Caloptilia* species (*C. azaleella* (Brants, 1913) and *C. leucothoes* Kumata, 1982) that develop on various *Rhododendron* L. species in Japan. In *C. azaleella*, which has two generations per year, dominant parasitoids included *Apanteles* cf. *xanthostigma*, *Pholetesor laetus* (Marshall, 1885) (Braconidae), *Acrolyta* sp. (Ichneumonidae), and *Sympiesis dolichogaster* (Eulophidae). For *C. leucothoes*, Sugiura (2011) reported the predominance of *Achrysocharoides* sp. (Eulophidae). Additionally, *S. dolichogaster* and other *Sympiesis* species were among the most frequently recorded parasitoids of *C. azaleella* in Mizell and Schiffhauer’s (1991) study conducted in the United States.

The number of parasitoid species recorded for *C. azaleella* (18 species) in Sugiura (2011) is comparable to the 15 species identified for *C. fidella* in our study, highlighting a similar degree of parasitoid diversity. However, *C. leucothoes*, which has only one generation per year, had a lower number of recorded parasitoid species (7). A comparable parasitoid composition was observed in Barry et al. (2010), who investigated *C. porphyretica* (Braun, 1923) in the United States. That study identified *Pholetesor* sp. (Braconidae) and Eulophidae as the most abundant parasitoids. Moreover, Wist and Evenden (2013), in their study on *C. fraxinella* (Ely, 1915) in Canada, noted that the dominant parasitoids were *Apanteles polychrosidis* Viereck, 1912 (Braconidae) and *Sympiesis sericeicornis*, which was also identified in this study alongside with two unidentified *Sympiesis* species.

In contrast to the dominant role of Braconidae found in many studies, Wheeler et al. (2017) reported that the parasitoid fauna of *C. triadicae* Davis, 2013 in the United States was dominated by *Goniozus* sp. (Bethylidae) and *Brasema* sp. (Eupelmidae). However, members of *Sympiesis* (Eulophidae) were also among the main parasitoids in that study. The differences in parasitoid composition for *C. triadicae*, an introduced species in the United States, were attributed to its non-native origin and the initial reliance on generalist parasitoids in the new environment. Over time, the parasitoid complex may shift as local parasitoids adapt to this new host, a phenomenon also observed by Grabenweger et al. (2010).

### Parasitism and Host Strategies

Leaf-mining lifestyles occur in the orders Lepidoptera, Diptera, Coleoptera, and Hymenoptera, and is diverse and abundant in Lepidoptera (Rott & Godfray, 2000, Kirichenko et al., 2018). Our results indicate that mining is a more effective strategy for reducing parasitism compared to leaf rolling, as *C. zieglerella*, which exclusively develops as a leaf-miner, and the mining stages of *C. fidella* exhibited lower parasitism rates than its leaf-rolling stages. Previous studies suggest that mining complexity deters parasitoids, making it harder for them to locate their hosts (Ayabe et al., 2008). For example, *Phyllonorycter malella* creates intricate mining networks that reduce parasitism risk by acting as barriers (Djemai et al., 2000). While our study species do not exhibit such extreme modifications, leaf mines (concealed stage) generally provide confined spaces where larvae can actively dislodge parasitoids. However, the relationship between host concealment and parasitism risk remains complex. Kobayashi et al. (2015) found that leaf-rolling weevils that seal or reinforce their leaf rolls (closed and wrapped rollers=concealed) experience lower parasitism rates than those that leave openings (open rollers=semi-concealed). Similarly, stem-boring (concealed) species exhibited lower parasitism rates than leaf-mining ones, suggesting that physical barriers can enhance protection. Similarly, Hrček et al. (2013) found that semi-concealed hosts were more frequently parasitized than exposed hosts, likely due to visual cues associated with leaf damage that parasitoids use to locate their hosts, the active defense of exposed hosts, or competition between parasitoid guilds. In our study, we classify leaf-mining stages as concealed hosts and leaf-rolling stages as semi-concealed hosts.

A key distinction in parasitoid communities is the balance between koinobionts and idiobionts. Koinobionts allow their hosts to continue development after parasitism, whereas idiobionts immediately immobilize and consume their hosts. Hrček et al. (2013) found that semi-concealed hosts were primarily parasitized by koinobionts, with Braconidae dominating, and exposed hosts were mostly attacked by koinobiont Tachinidae. These two host feeding-mode groups also did not differ in parasitoid community size. In contrast, leaf-mining species are known to be frequently parasitized by Chalcidoidea and Ichneumonidae, with a higher prevalence of idiobionts (Askew & Shaw, 1979, Hawkins, 1994). Our study contrasts with Hrček et al. (2013), as we recorded a higher proportion of idiobionts in both concealed leaf-mining and semi-concealed leaf-rolling stages, particularly in *C. fidella*. This might suggest that the koinobiont-idiobiont ratio varies significantly depending on host species, habitat, and parasitoid community composition.

In this context, the semi-concealed stage of *C. fidella* provides some protection but remains more vulnerable to parasitoids than the fully concealed leaf-mining stage. Larvae in rolls either bend the tip of a leaf or fold its margin, creating a pocket-like shelter that never fully seals, making it more susceptible to parasitoids than species that construct tightly sealed shelters (Kobayashi et al., 2015). This may explain why its leaf-rolling stages had higher parasitism rates than its mining stages, despite the additional structural protection.

Overall, our findings support the hypothesis that semi-concealed hosts experience the highest level of parasitism risk between fully exposed and concealed species as in Hrček et al. (2013). However, such results might be influenced by host concealment type, parasitoid searching strategies, and the koinobiont-idiobiont balance. Early gracillariids likely fed on Fabales tree leaves, forming simple blotch mines, while later adaptations, such as keeled or tentiform blotch mines, leaf rolling, and gall formation, emerged as defensive strategies (Li et al., 2022; Aoyama & Ohshima, 2019). However, in contrast to various mining strategies, leaf rolling evolved as the most advanced larval behaviour mode only in a few clades of Gracillariidae (incl. Gracillariinae, subf. of *Calloptilia*) (Li et al., 2022, Kawahara et al., 2017). Although Li et al. (2022) suggested advanced larval behaviours, such as making keeled or tentiform blotch mines, rolling leaves and making galls, may have accelerated the diversification of Gracillariidae by avoiding larval parasitoids, our results indicate leaf rolling might not be a sufficient strategy to avoid parasitism at least in *C. fidella*. Further studies with expanding taxonomic sampling across gracillariid hosts could provide further insights into how parasitoid-host interactions shape the evolution of leaf-mining and leaf-rolling behaviors in Gracillariidae.

## CONCLUSION

This study highlights how the level of host concealment influences parasitoid communities and parasitism rates in two microlepidopteran species on hops. We demonstrate that *C. fidella*, which transitions from mining to leaf rolling, is more vulnerable to parasitoids in its semi-concealed stage than in its mining stage. In contrast, *C. zieglerella*, which remains a leaf miner throughout its development, experiences lower parasitism rates, supporting the idea that mining is a more effective strategy for reducing parasitism risk. The parasitoid assemblages associated with the two hosts differ, with *C. fidella* hosting a diverse range of parasitoids, including braconids and eulophids, and *C. zieglerella* being predominantly parasitized by eulophids. The higher proportion of idiobiont parasitoids in leaf rolls suggests that this developmental stage is more susceptible to parasitoidism by more generalist idiobionts.

Beyond ecological insights, this study demonstrates the effectiveness of molecular species delimitation in resolving taxonomic uncertainties within parasitoid communities. The combination of ITS2 and CO1 markers, applied using the bPTP and ASAP methods, proved valuable in identifying species and detecting cryptic diversity. While both methods provided useful results, bPTP was more sensitive in delineating putative cryptic species, whereas ASAP yielded a more conservative classification. These findings emphasize the necessity of integrating molecular tools with morphological and ecological data to achieve robust species identification.

Furthermore, the molecular data confirmed the idiobiont strategy in *S. acalle*, *S. sericeicornis*, and *E. fenestratus*, providing new insights into the functional roles of parasitoids associated with microlepidopteran hosts. This study underscores the importance of an integrative taxonomic approach for understanding host-parasitoid interactions, as molecular tools can refine species identification and reveal hidden diversity within parasitoid assemblages.

Future research should expand the taxonomic scope of host-parasitoid systems to assess how these interactions shape the evolution of feeding and sheltering behaviors in phytophagous insects. Additionally, further exploration of cryptic diversity within parasitoid communities will enhance our understanding of their ecological roles and evolutionary dynamics.

## ACKNOWLEDGEMENTS

This project was supported by the Grant Agency of Charles University (GAUK), project no. 375421 (TH) and grant PRIMUS/24/SCI/015 (PJ). We are grateful to our colleagues at the Department of Zoology, Faculty of Science, Charles University; the Department of Entomology, National Museum of the Czech Republic (institutional support project IP DRKVO 2024–2028/5.I.a, EORI 00023272); the Department of Entomology, State Museum of Natural History Stuttgart; and the Department of Integrated Crop Protection against Pests, Czech Agrifood Research Center (institutional support MZE-RO0423) and to Nela Gloríková, for their valuable collaboration and support. We also thank Rudolf Bryner for providing the photograph of an adult *Cosmopterix zieglerella* (Figure 3H).

## SUPPLEMENTARY FILES

**Figure S1.**
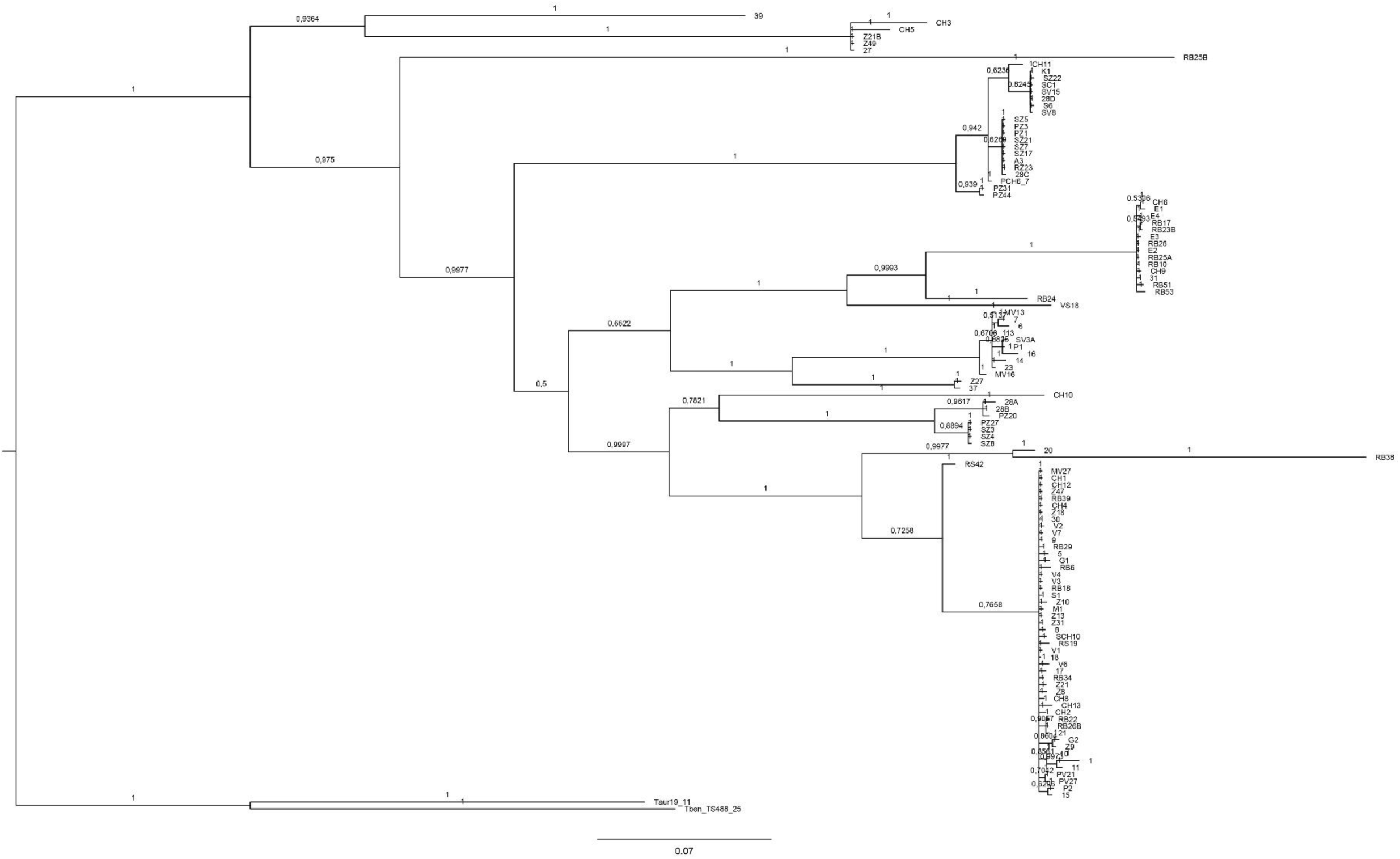
Phylogenetic tree of Chalcidoidea parasitoids based on MrBayese analysis of concatenated COI and ITS2 sequences.

**Figure S2.**
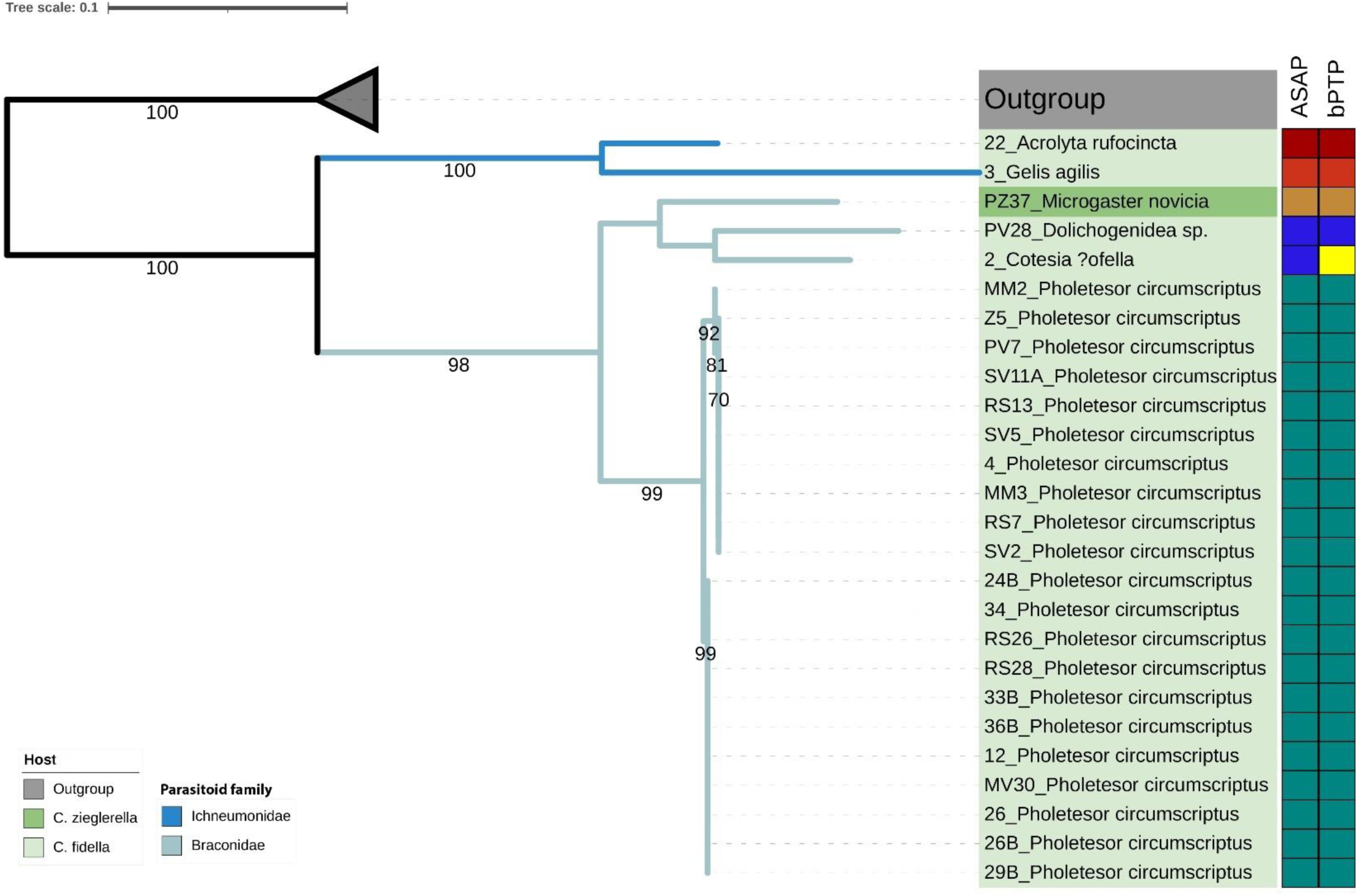
Phylogenetic tree of Ichneumonoidea parasitoids based on RAxML analysis of COI sequences. Bootstrap values >70 are indicated at branch nodes. Branch colors correspond to different families, while the background shading of parasitoid species represents their respective hosts. Additionally, colors distinguish the delimitation based on genetic distances using the ASAP and bPTP methods.

**Figure S3.**
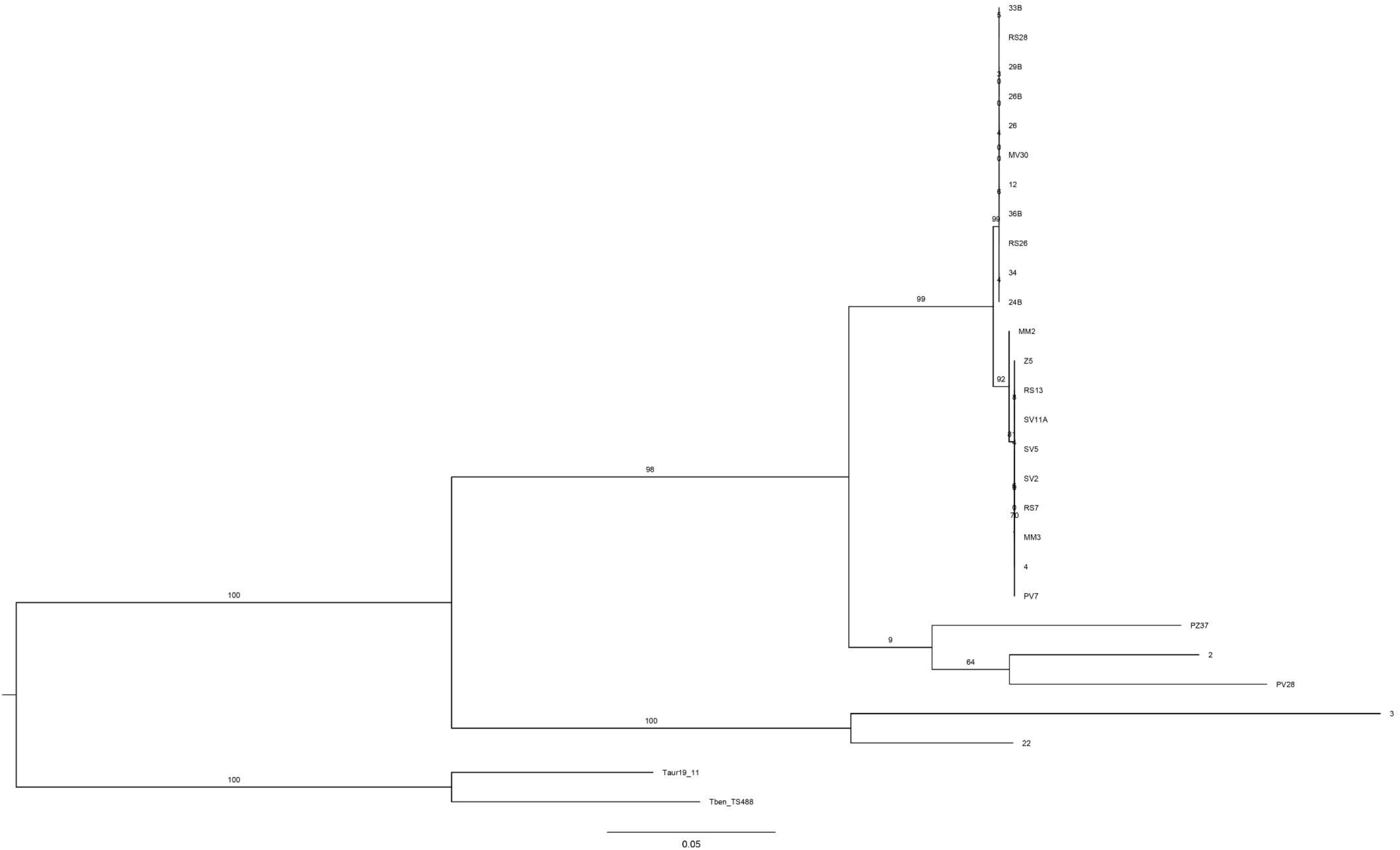
Phylogenetic tree of Ichneumonoidea parasitoids based on RAxML analysis of COI.

**Figure S4.**
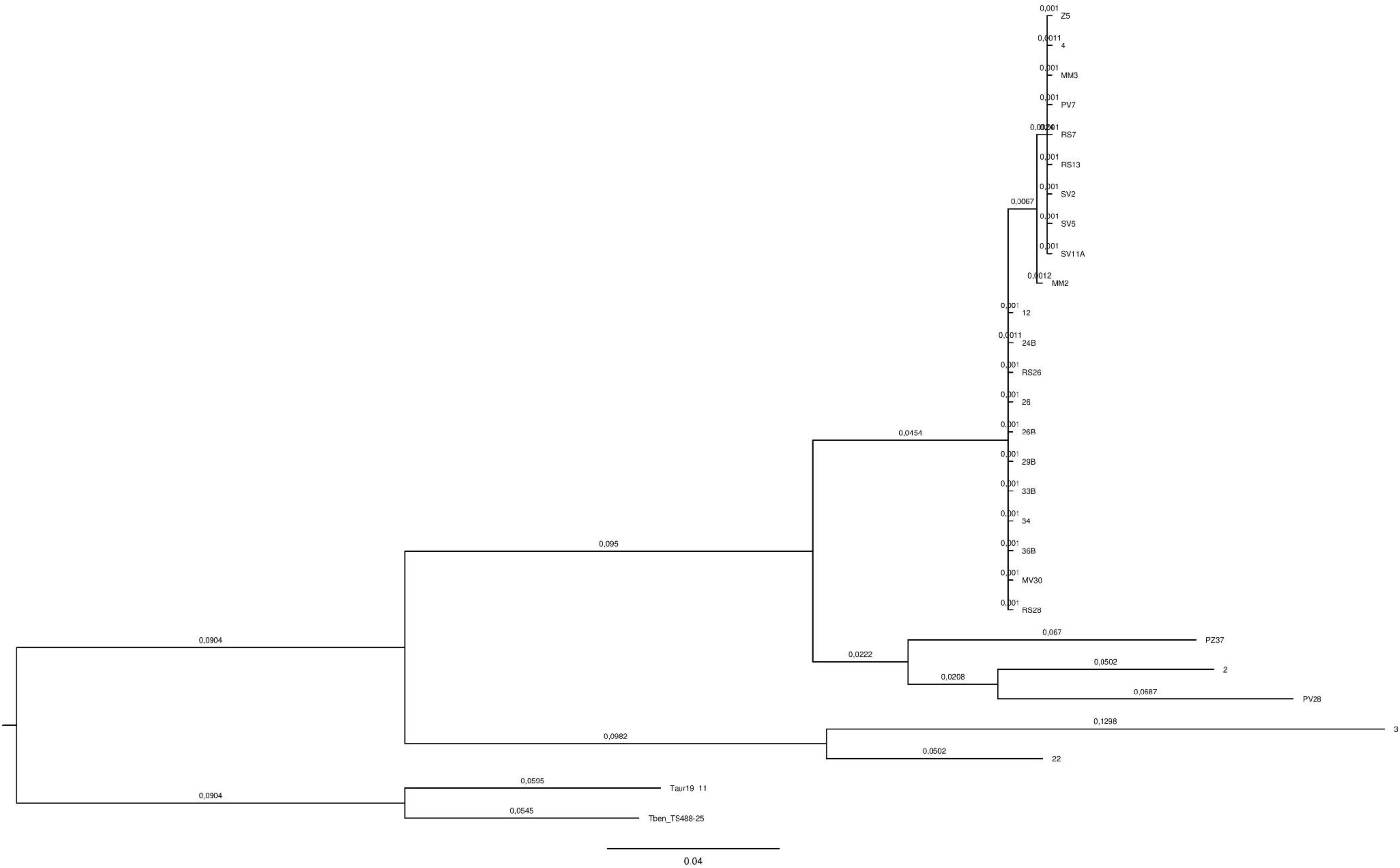
Phylogenetic tree of Ichneumonoidea parasitoids based on MrBayese analysis of COI.

